# GNL3 SUMOylation is essential for DNA double-strand break repair by homologous recombination

**DOI:** 10.1101/2025.11.04.686352

**Authors:** Yunhan Yang, Yanping Li, Roselyn S. Dai, Canping Chen, Keith Zientek, Ashok Reddy, Zheng Xia, Rosalie C. Sears, Xiao-Xin Sun, Mu-Shui Dai

**Affiliations:** Department of Molecular & Medical Genetics, School of Medicine, Oregon Health & Science University, 3181 SW Sam Jackson Park Road, Portland, OR 97239, USA; Department of Biomedical Engineering, Oregon Health & Science University, 3181 SW Sam Jackson Park Road, Portland, OR 97239, USA; OHSU Proteomics Shared Resource, Oregon Health & Science University, 3181 SW Sam Jackson Park Road, Portland, OR 97239, USA; Center for Biomedical Data Science, Oregon Health & Science University, 3181 SW Sam Jackson Park Road, Portland, OR 97239, USA; the OHSU Knight Cancer Institute, Oregon Health & Science University, 3181 SW Sam Jackson Park Road, Portland, OR 97239, USA; Brenden-Colson Center for Pancreatic Care, Oregon Health & Science University, 3181 SW Sam Jackson Park Road, Portland, OR 97239, USA

## Abstract

DNA double-strand break (DSB) repair via homologous recombination (HR) is critical for maintaining genomic integrity and requires proper DNA end resection to generate single-stranded DNA (ssDNA) overhangs. However, the mechanisms governing this critical step in cells remains poorly understood. Here, we report that GNL3, a nucleolar GTP-binding protein, plays a key role in HR repair via regulating DNA end resection dependently on its SUMOylation. Ectopic expression of wild-type, but not the SUMO-defective K196R mutant, GNL3 completely abolished DNA damage response induced by knockdown of endogenous GNL3. GNL3 interacts with the BLM-DNA2 helicase-nuclease complex and is critical for DNA end resection and subsequent loading of RPA and RAD51. This interaction requires SUMOylation and SUMO-interacting motifs (SIMs) in both proteins. We further demonstrate that USP36, a nucleolar deubiquitinating enzyme, functions as a novel SUMO ligase for GNL3, while the SUMO protease SENP3 deSUMOylates GNL3. Notably, several breast cancer-derived GNL3 variants that disrupt its SUMOylation or SIM fail to interact with the BLM-DNA2 complex. Knockdown of GNL3 sensitizes HR-proficient breast cancer cells to etoposide or Olaparib treatment. Together, our results reveal that GNL3 SUMOylation is crucial for HR repair and suggest that targeting GNL3 SUMOylation may induce HR deficiency, thereby sensitizing breast cancers to DNA damage-inducing agents.

## Introduction

DNA double-strand break (DSB) induced by both endogenous and exogenous DNA damage agents such as replication stress, irradiation (IR), and toxic chemicals represents the most detrimental form of genomic lesions in cells ^1^. Cells develop a fundamental and sophisticated process called DNA damage response (DDR) to sense and repair DNA breaks to ensure genomic integrity ^1,2^. Eukaryotic cells utilize two major pathways to repair DSBs: non-homologous end joining (NHEJ) directly rejoins the DSB ends but is error-prone ^3^, whereas homologous recombination (HR) provides high-fidelity repair by using a sister chromatid as the template during the S and G2 phases of the cell cycle and is also the primary mechanism for repairing replication-associated DSBs ^4,5^. A third backup mechanism called microhomology-mediated end-joining (MMEJ) repairs DSBs by DNA polymerase theta (Polθ) during mitosis or when HR and NHEJ are compromised and is inherently mutagenic^6^. Unrepaired or incorrectly repaired DSBs lead to genomic instability, cell death and various diseases including cancer ^1^.

DNA end resection is an essential process in HR and determines the choice of HR over NHEJ ^5,7,8^. The MRE11-RAD50-NBS1 (MRN) complex senses DSBs and recruits the ATM kinase to phosphorylate downstream effector proteins to amplify and orchestrate the cellular response to DSBs ^8–10^. MRE11 initiates DNA end resection by cleaving the 5’ strand of the DSB ends with the cooperation of its phosphorylated CtIP cofactor, generating a short stretch of 3’ single-stranded DNA (ssDNA), which serves as a docking site for the long-range resection exonuclease 1 (Exo1) or the DNA helicase BLM and the DNA replication helicase/nuclease 2 (DNA2) complex ^7,8^. The helicases and nucleases work to unwind the DNA double helix and digest the 5’ DNA strand to generate a long stretch of 3’ ssDNA^7,8^. The resulting ssDNA is rapidly coated and stabilized by the replication protein A (RPA) complex, which is then replaced by the RAD51 recombinase to form a nucleofilament, a step that requires the BRCA1-PALB2-BRCA2 complex ^5,10,11^. The RAD51-ssDNA nucleoprotein filaments conduct a homology search and catalyze DNA strand exchange for high-fidelity repair of DSBs by HR^12^.

While defects in HR pathway result in genomic instability and tumorigenesis, cancers with HR deficiency are synthetic lethal to treatment with poly(ADP-Ribose) polymerase inhibitors (PARPi), including tumors with BRCA1 mutations or with somatic mutations in other DDR and repair pathway genes in the absence of BRCA1 mutations (called “BRCAness”)^13^. Yet, the majority of cancers are HR-proficient and PARPi resistant. HR-deficient cancers, although initially effective, often become resistant to PARPi due to BRCA1/2 reversion mutations or other adaptive mechanisms that restore the HR pathway^14–16^. Thus, there is an unmet need to identify new HR pathway targets to clinically re-sensitize cancers to PARPi treatment.

The DDR pathways are tightly regulated by post-translational modifications, including SUMOylation, a post-translational modification of proteins by <u>s</u>mall <u>u</u>biquitin-like <u>mo</u>difiers (SUMOs). Mammals express three main SUMO isoforms: SUMO2 and SUMO3 are 97% identical and each shares 45% sequence identity with SUMO1 ^17,18^. SUMOylation is ATP-dependent and occurs through sequential reactions involving a heterodimeric SUMO-activating enzyme SAE1/SAE2 (E1), a single SUMO-conjugating enzyme Ubc9 (E2) and one of a few SUMO ligases (E3) ^17,18^. The SUMO acceptor lysine (Lys, K) is often present within a conserved ΨKxE motif, where Ψ is a large hydrophobic amino acid and x is any amino acid ^19^. Various DDR pathway proteins^20–23^ are regulated by SUMOylation and SUMOylated proteins are enriched at local sites of DNA damage^24,25^. For example, SUMOylation of MDC1 upon DNA damage is critical for HR by promoting its RNF4-mediated ubiquitination, degradation and removal from DNA damage sites^26–29^. MRE11 requires SUMOylation to prevent its ubiquitination and degradation and function in DNA end resection ^30^. BLM SUMOylation at K317 and K331 is necessary for its interaction with RAD51, RAD51 accumulation at stalled DNA forks^31^, and HR repair^32,33^. Other DDR proteins subjected to SUMO regulation include BRCA1^27–29^, BARD1^34,35^, RAD18^35^, RPA1^36^, RAD51^37–39^, RAD52^40–43^, Ku70^44^, and the SUMO-targeted ubiquitin ligase (STUBL) RNF4 ^27–29,45^, playing roles in both HR and NHEJ^27–29,34,46–50^.

GNL3, also called nucleostemin, is a nucleolar GTP-binding protein originally identified in neuronal stem cells essential for stemness^51^. Earlier work showed that GNL3 plays a role in ribosome biogenesis^52,53^, p53 signaling and cell cycle regulation^54–57^. High expression of GNL3 has been observed in various cancers and cancer stem cells^58–62^. Interestingly, GNL3 is implicated in replication stress-induced DNA damage and HR repair^63–65^. However, how GNL3 is regulated in cells and how it is involved in HR repair are unclear.

Here, we report that GNL3 is regulated by SUMOylation and this SUMOylation plays a key role in HR repair. DNA damage promotes GNL3 SUMOylation and its interaction with the BLM-DNA2 complex. We showed that GNL3 contains a SUMO-interacting motif (SIM) and its interaction with BLM requires SUMO-SIM interactions between the two proteins. We further discovered that GNL3 is SUMOylated by USP36, a nucleolar deubiquitinase and SUMO ligase dual-function enzyme^66–70^, and deSUMOylated by the SUMO protease SENP3. Several breast cancer-derived GNL3 variants abolish its SUMO regulation in response to DSBs. Knockdown of GNL3 sensitizes HR-proficient TNBC cells to genotoxic agents and PARPi. Thus, USP36 and SENP3-regulated GNL3 SUMOylation is critical for DNA end resection during HR repair and thus GNL3 and its SUMOylation may represent important cancer therapeutic targets.

## Results

### GNL3 is SUMOylated in cells in response to DNA damage

To understand how GNL3’s role in HR is regulated, we sought to test if GNL3 is regulated by SUMOylation, given that SUMOylation plays a critical role in DDR ^23,71^. Indeed, GNL3 can be modified by both SUMO1 and SUMO2 (Fig. 1a & Supplementary Fig. 1a) and overexpression of Ubc9 markedly increased GNL3 SUMOylation, whereas dominant-negative Ubc9 mutant suppressed GNL3 SUMOylation in cells (Supplementary Fig. 1b). We next examined whether GNL3 can be SUMOylated in response to DNA damage. We found that treatment of etoposide (ETO), a topoisomerase 2 inhibitor that induces DSBs^72,73^, markedly induced GNL3 SUMOylation by SUMO2 in U2OS (Fig. 1b) and HeLa (Fig. 1c) cells, peaking at 2 hours following the ETO treatment. GNL3 SUMOylation induced by ETO treatment is completely blocked by the SUMO activating enzyme (E1) inhibitor ML792 (Fig. 1d). Interestingly, GNL3 SUMOylation by SUMO1 remained unchanged upon ETO treatment (Fig. 1e). GNL3 SUMOylation by SUMO2 was also increased in cells treated with the topoisomerase 1 inhibitor camptothecin (CPT) and the replication stress inducer hydroxyurea (HU) (Fig. 1f). These data reveal that DNA damage promotes GNL3 SUMOylation by SUMO2. Chromatin fractionation assays showed that chromatin-associated GNL3 is significantly increased upon ETO treatment (Fig. 1g) and GNL3 SUMOylation in the chromatin fraction is significantly induced by the ETO treatment (Fig. 1h). Together, these results suggest that GNL3 modification by SUMO2 on chromatin may play a role in DNA damage response and repair.

**Figure 1.**
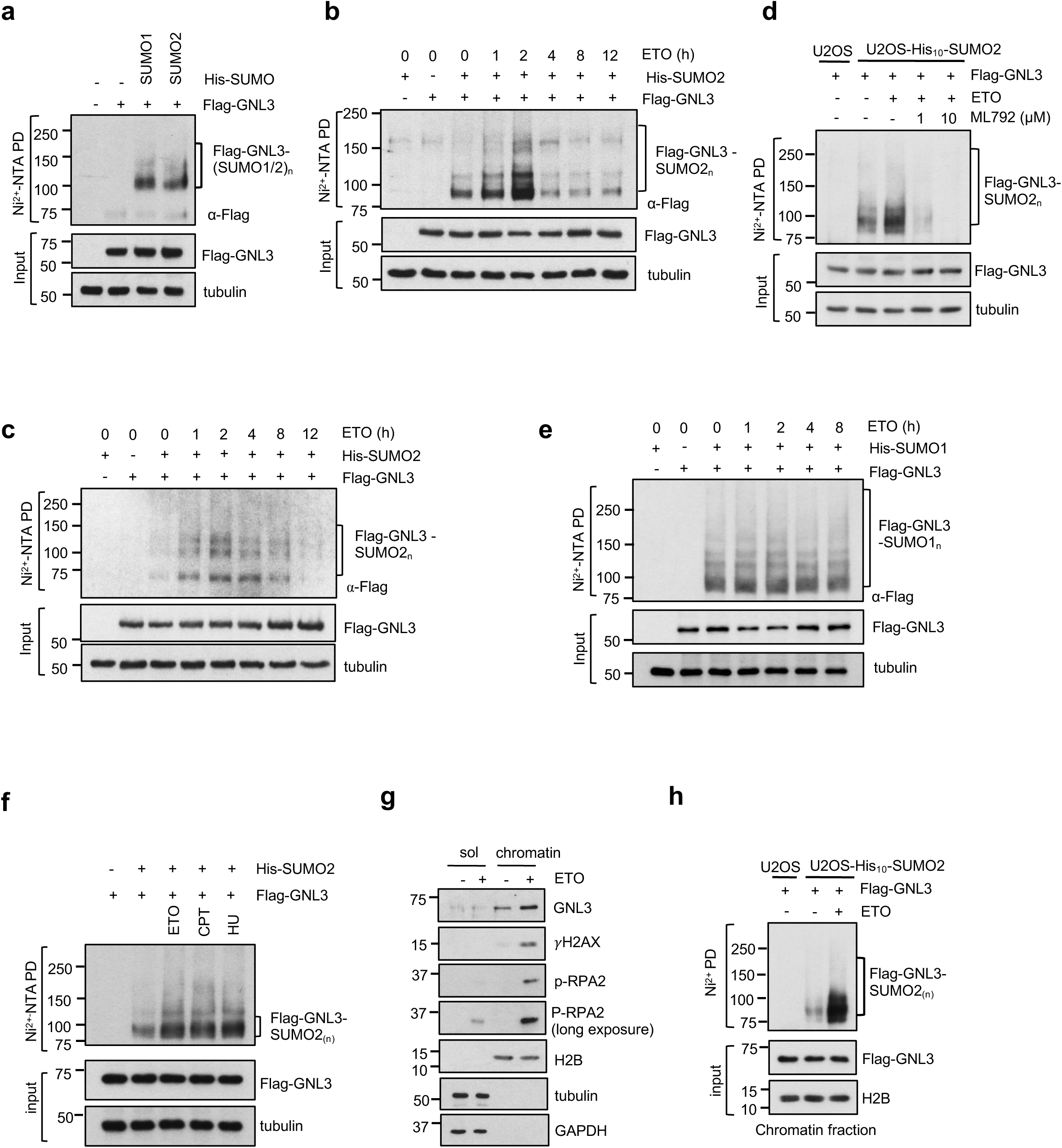
DNA damage induces GNL3 SUMOylation. **(a)** GNL3 is modified by both SUMO1 and SUMO2. H1299 cells transfected with the indicated plasmids were subjected to Ni^2+^-NTA pulldown (PD) under denaturing conditions, followed by IB with anti-Flag antibody. **(b)(c)** GNL3 SUMOylation by SUMO2 is increased in response to DNA damage. U2OS (B) and HeLa (C) cells transfected with the indicated plasmids were treated with 20 μM ETO for the indicated times or control DMSO and assayed for GNL3 SUMOylation by Ni^2+^-NTA pull-down (PD) under denaturing conditions followed by IB. The SUMO2-modified GNL3 is indicated. The protein expression is shown in the bottom panels. **(d)** DNA damage-induced GNL3 SUMOylation is abolished by SUMO E1 inhibitor treatment. His-SUMO2 stably expressing U2OS cells were treated with 20 μM ETO in the presence of the indicated doses of ML792 and assayed by IB. **(e)** DNA damage does not promote GNL3 SUMOylation by SUMO1. U2OS cells transfected with the indicated plasmids were treated with DMSO or with 20 μM ETO for the indicated times and assayed for GNL3 SUMOylation by Ni^2+^-NTA PD. The SUMO1-modified GNL3 is indicated. **(f)** GNL3 SUMOylation is increased in response to DNA damage agents. U2OS cells stably expressing His_10_-SUMO2 were treated with control DMSO, 20 μM ETO or 2 μM CPT for 2 hours or 10 mM HU for 4 hours and then assayed for GNL3 SUMOylation. **(g)** GNL3 associates with chromatin in response to DNA damage. U2OS cells were fractionated into soluble and chromatin fractions and assayed by IB. **(h)** GNL3 SUMOylated on chromatin in response to DSBs. U2OS and U2OS-His_10_-SUMO2 cells transfected with Flag-GNL3 plasmid were treated with DMSO or 20 μM ETO for 2 hours. Chromatin fractions were isolated and assayed for GNL3 SUMOylation by Ni^2+^-NTA PD under denaturing conditions followed by IB.

### GNL3 SUMOylation at K196 is required for DSB repair and cell viability

To define the SUMOylation sites in GNL3 protein, we mutated the two consensus SUMO lysine (K) residues predicted by the GPS-SUMO tool ^74^, K196 and K275 (Fig. 2a), to Arg (R) and examined their SUMOylation in cells. As shown in Fig. 2b, mutating K196, but not K275, largely abolished GNL3 SUMOylation, suggesting that GNL3 is SUMOylated at K196. To examine whether GNL3 SUMOylation is critical for its role in HR, we performed knockdown and rescue experiments. Consistent with previous studies ^61,63^, knockdown of GNL3 markedly induced DDR as shown by the increased levels and foci formation of phosphorylated H2AX (γH2AX) in U2OS cells (Figs. 2c-2e). Interestingly, ectopic expression of wild-type (WT) GNL3, but not the K196R mutant, abolished the induction of γH2AX foci (Figs. 2c & 2d) and levels (Fig. 2e) induced by knockdown of endogenous GNL3, but not in response to ETO treatment (Supplementary Figs. 2a & 2b). WT GNL3, but not the K196R mutant, also abolished the induction of γH2AX as well as phosphorylated RPA2 induced by knockdown of endogenous GNL3 in breast cancer MDA-MB-231 cells (Figs. 2f & 2g). Consistently, ectopic expression of WT GNL3, but not the K196R mutant, abolished the induction of p53 levels in response to DNA damage induced by knockdown of endogenous GNL3 (Supplementary Fig. 2e). Additionally, knockdown of GNL3 markedly inhibited DSB repair that can be rescued by ectopic expression of WT GNL3, but not the K196R mutant, as determined by Comet assays (Figs. 2h & 2i), but not in the presence of ETO treatment (Supplementary Figs. 2c & 2d). Cell viability (Fig. 2j) and colony formation (Figs. 2k & 2l) assays reveal that knockdown of GNL3 significantly inhibited U2OS cell growth, which is rescued by expression of WT GNL3, but not the K196R mutant. Together, these results suggest that GNL3 SUMOylation is critical for DSB repair and cell viability.

**Figure 2.**
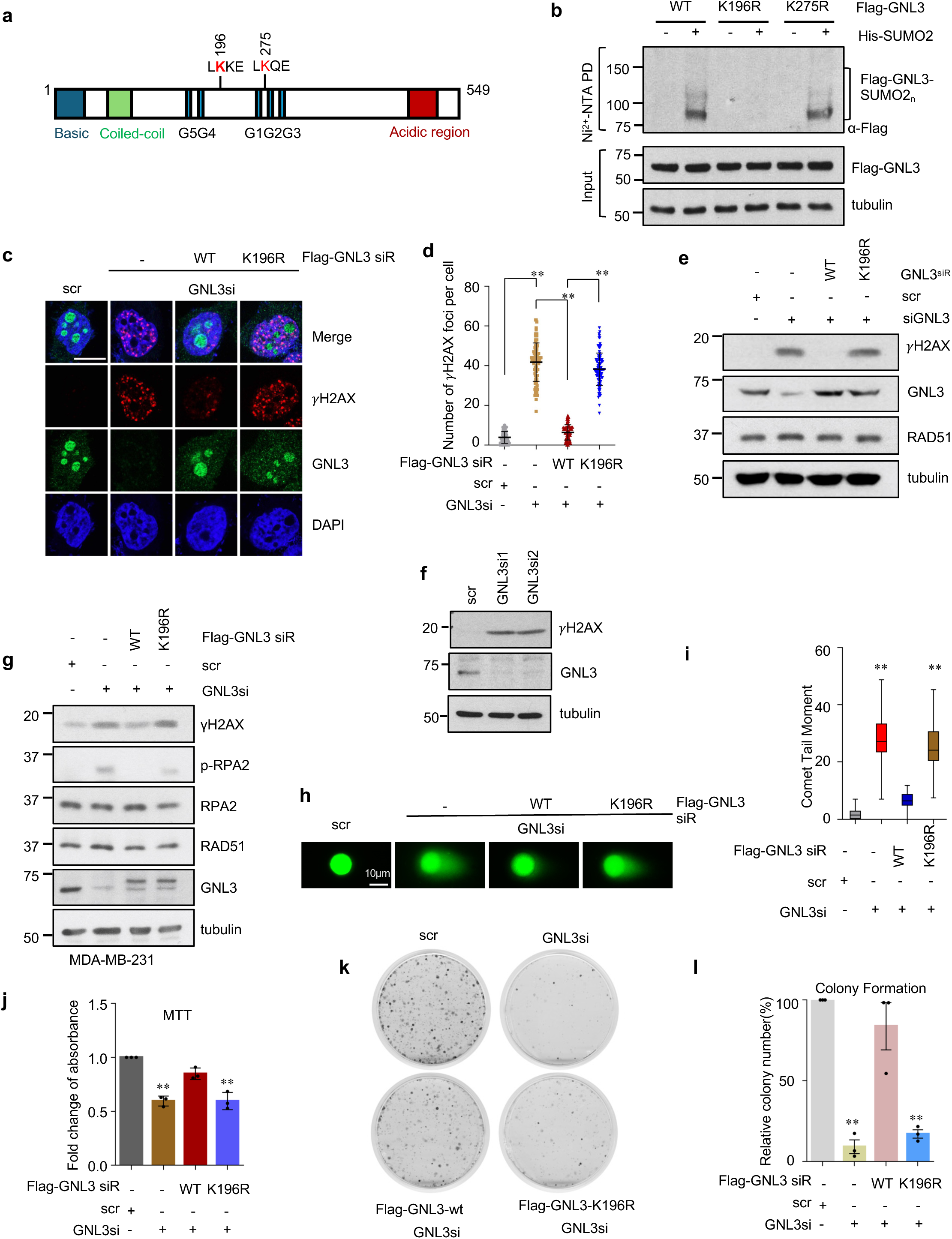
GNL3 SUMOylation at K196 is essential for DDR. **(a)** Diagram of GNL3 protein with the indication of the putative SUMO sites and its associated basic, coiled-coil, G1-G5 GTP-binding motifs, and acidic domain. **(b)** GNL3 is SUMOylated at K196. H1299 cells transfected with WT GNL3 or the indicated lysine mutant plasmids without or with His-SUMO2 were subjected to Ni^2+^-NTA PD under denaturing conditions followed by IB. The SUMOylated GNL3 is indicated. The protein expression is shown in the bottom panels. **(c)(d)** Ectopic expression of WT GNL3, but not the K196R mutant, abolishes γH2AX foci formation upon knockdown of endogenous GNL3. U2OS cells transfected with scrambled (scr) or GNL3 siRNA together with Flag-tagged siRNA-resistant (siR) WT GNL3 or the K196R mutant were assayed by IF staining with anti-GNL3 and anti-γH2AX antibodies. Shown are representative confocal images (c) and the quantification (d). **(e).** Ectopic expression of WT GNL3, but not the K196R mutant, abolishes GNL3 knockdown-induced H2AX phosphorylation. U2OS cells transfected with scr or GNL3 siRNA together with siRNA-resistant WT GNL3 or the K196R mutant were assayed by IB. **(f)** Knockdown of GNL3 induces the levels of γH2AX in MDA-MB-231 cells. **(g)** Ectopic expression of WT GNL3, but not the K196R mutant, abolishes GNL3 knockdown-induced γH2AX. MDA-MB-231 cells transfected with scr or GNL3 siRNA together with siRNA-resistant WT Flag-GNL3 or the K196R mutant were assayed by IB. **(h)(i)** GNL3 SUMOylation is essential for DSB repair. U2OS cells transfected with scr or GNL3 siRNA together with Flag-tagged siRNA-resistant WT GNL3 or the K196R mutant were subjected to Comet assays. Shown are representative images (h) and the quantification (i). **(j)-(l)** GNL3 SUMOylation is essential for cell proliferation. U2OS cells transfected with scr or GNL3 siRNA together with Flag-tagged siRNA-resistant WT GNL3 or the K196R mutant were assayed by MTT cell viability (j) and colony formation assays (k)(l). Representative images (k) and the quantification (l) of the colony formation are shown. Scale bar, 10 μm. **P<0.01, compared to scr control.

### GNL3 SUMOylation is critical for its interaction with γH2AX and RAD51

To determine whether GNL3 localizes to DNA damage sites, we examined whether it interacts with γH2AX, an early DNA damage marker responsible for recruiting DDR proteins to damage sites. We performed proximity ligation assays (PLAs) and observed that treatment of cells with either ETO or X-rays markedly induced the interaction between GNL3 and γH2AX (Figs. 3a & 3b). The interaction was also shown by using co-immunoprecipitation (co-IP) assays, peaking at 2 hours after ETO treatment (Fig. 3c). Interestingly, abolishing GNL3 SUMOylation by mutating K196 markedly reduced its interaction with γH2AX (Fig. 3c). These results suggest that GNL3 is recruited to the DNA damage site dependently on its SUMOylation. It has previously been shown that GNL3 interacts with RAD51, the recombinase essential for HR repair ^63^. Indeed, PLA assays showed that the interaction between GNL3 and RAD51 is also markedly increased following the treatment with either X-rays or ETO (Figs. 3d & 3e). Co-IP assays also showed that the GNL3-RAD51 interaction peaked at 2 hours after ETO treatment, which is markedly impaired by mutating K196 (Fig. 3c). Together, these data suggest that GNL3 SUMOylation is critical for its recruitment to chromatin and interaction with RAD51.

**Figure 3.**
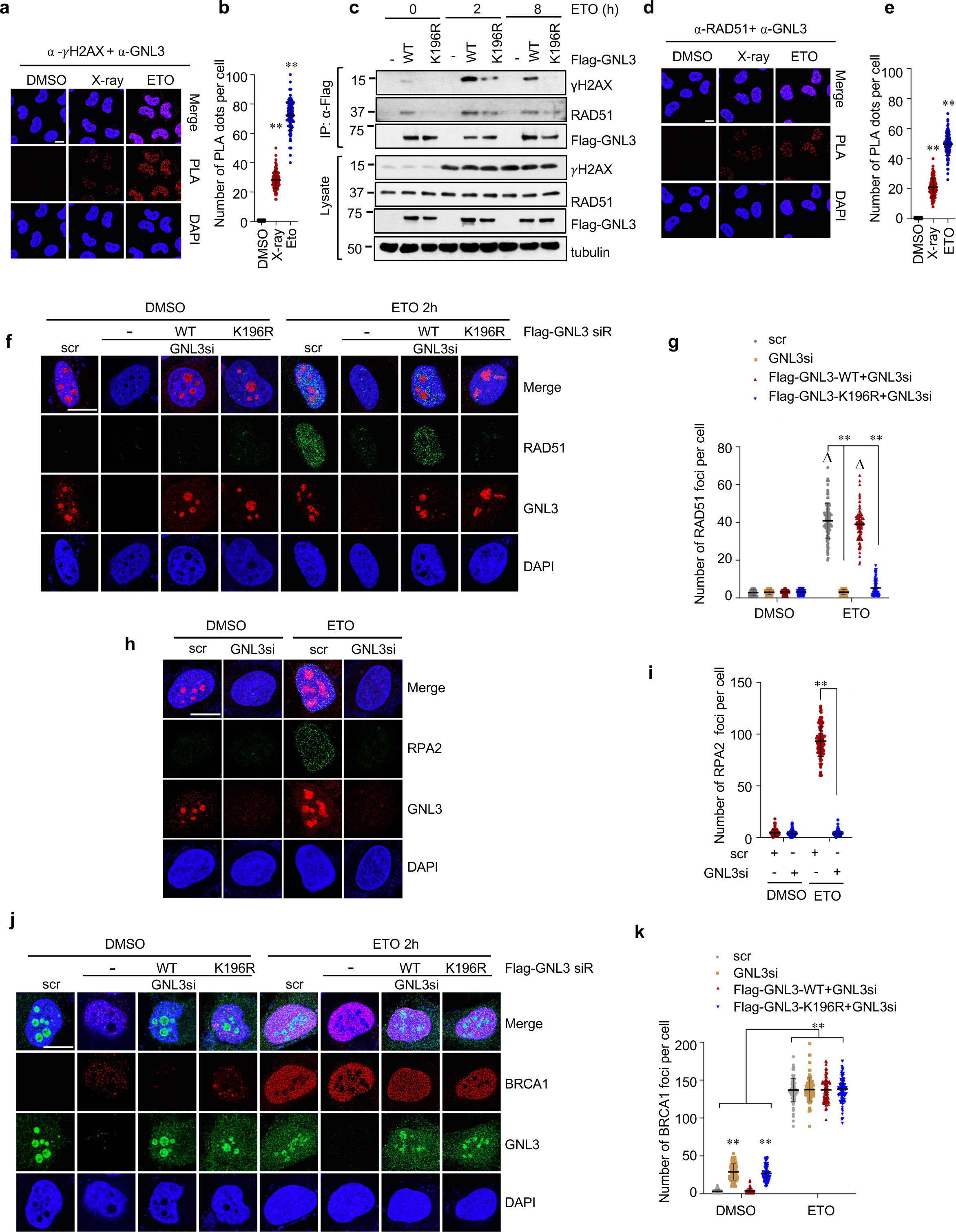
GNL3 SUMOylation is required for interaction with RAD51 and γH2AX and the RAD51 and RPA foci formation in response to DSBs. **(a)(b)** DNA damage induces GNL3 interaction with γH2AX. U2OS cells treated with 10 Gy X-ray irradiation or 20 μM ETO for 2 hours were assayed by PLA using anti-GNL3 and anti-γH2AX antibodies. Shown are representative confocal images (a) and the quantification (b). **(c)** GNL3 SUMOylation is critical for GNL3 interaction with γH2AX and RAD51 in response to DNA damage. U2OS cells transfected with indicated plasmids were treated with 20 μM ETO for the indicated times were subjected to IP using anti-Flag antibody followed by IB. **(d)(e)** DNA damage induces GNL3 interaction with RAD51. U2OS cells treated with 10 Gy X-ray irradiation or 20 μM ETO for 2 hours were assayed by PLA using anti-GNL3 and anti-RAD51 antibodies. Shown are representative confocal images (d) and the quantification (e). **(f)(g)** GNL3 SUMOylation is critical for RAD51 foci formation upon DNA damage. U2OS cells transfected with scr or GNL3 siRNA together with Flag-tagged and siRNA-resistant WT GNL3 or the K196R mutant were treated without or with 20 μM ETO and assayed for RAD51 foci formation by IF with anti-RAD51 and anti-GNL3 antibodies. Shown are representative confocal images (f) and quantification (g). **(h)(i)** GNL3 is also critical for RPA foci formation upon DNA damage. U2OS cells transfected with scr or GNL3 siRNA were treated without or with 20 μM ETO and assayed for RPA foci formation by IF with anti-RPA2 phospho-Ser33 and anti-GNL3 antibodies. Shown are representative confocal images (h) and quantification (i). **(j)(k)** GNL3 SUMOylation is not required for BRCA1 foci formation upon DNA damage. U2OS cells transfected with scr or GNL3 siRNA together with Flag-tagged and siRNA-resistant WT GNL3 or the K196R mutant were treated without or with 20 μM ETO and assayed for BRCA1 foci formation by IF with anti-BRCA1 and anti-GNL3 antibodies. Shown are representative confocal images (j) and quantification (k). Scale bar, 10 μm. **P<0.01, compared to the control group. △P<0.01, compared to the respective DMSO-treated group (triangle symbol).

### GNL3 SUMOylation is required for RAD51 and RPA2 foci formation in response to DSBs

To understand how SUMOylation regulates GNL3 function in HR repair, we examined the DNA damage-induced foci formation of key DDR proteins. We found that knockdown of GNL3 does not abolish γH2AX foci formation in cells treated with ETO (supplementary Fig. 2a & 2b). Interestingly, DNA damage induced by depleting endogenous GNL3 fails to induce RAD51 foci (Figs. 3f & 3g), whereas knockdown of GNL3 abolishes ETO treatment-induced RAD51 foci formation, which can be completely rescued by ectopic expression of WT GNL3, but not the K196R mutant (Figs. 3f & 3g). Knockdown of GNL3 also abolishes RPA2 foci formation induced by ETO treatment (Figs. 3h & 3i). Intriguingly, knockdown of GNL3 does not abolish the foci formation of BRCA1 (Figs. 3j, 3k), BARD1 (Supplementary Figs. 3a & 3b), 53BP1 (Supplementary Figs. 3c & 3d) and MDC1 (Supplementary Figs. 3c & 3e) following the ETO treatment. These results suggest that GNL3 may play a role in HR repair by regulating the RPA and RAD51 loading without interfering with the tested DDR proteins, including BRCA1, BARD1 MDC1 and 53BP1. Of note, SUMOylation does not affect the nucleolar localization of GNL3 as the K196R mutant still predominantly localized in the nucleolus, similar to WT GNL3, as determined by immunofluorescence staining, where the GTP-binding-defective G261V mutant is localized in the nucleoplasm (Supplementary Fig. 3f)

### GNL3 regulates DNA end resection by interacting with the BLM-DNA2 complex in response to DSBs dependently on its SUMOylation

As GNL3 SUMOylation is required for RAD51 and RPA foci formation and our affinity purification and mass spectrometry analyses identified BLM in the Flag-GNL3-associated protein complexes (Fig. 4a), we asked whether GNL3 SUMOylation may play a role in DNA end resection. We first test whether GNL3 interacts with the BLM-DNA2 complex critical for long-range DNA end resection. Indeed, co-IP assays showed that treatment of cells with ETO markedly promoted Flag-GNL3 interaction with endogenous BLM and DNA2 (Fig. 4b). PLA assays showed that GNL3 interacts with BLM endogenously upon ETO treatment (Figs. 4c & 4d). Flag-BLM also co-immunoprecipitated with endogenous GNL3 that is significantly induced by ETO treatment (Fig. 4e). Interestingly, the interaction of GNL3 with the BLM-DNA2 complex in response to ETO treatment was markedly attenuated by mutating the SUMOylation lysine K196 (Figs. 4b, 4f, 4g & Supplementary Fig. 4). We next examined whether GNL3 plays a role in DNA end resection. We performed 5-bromo-2’-deoxyuridine (BrdU) incorporation assays followed by immunofluorescence (IF) staining under native conditions with anti-BrdU antibody, which only recognizes ssDNA in non-denaturing conditions ^75^. As shown in Figs. 4h and 4i, knockdown of GNL3 markedly reduced the anti-BrdU staining positive cells upon ETO treatment, indicative of the impaired DNA end resection and ssDNA generation, which can be again rescued by WT GNL3, but not the K196R mutant. Thus, GNL3 SUMOylation plays a key role in regulating DNA end resection by interacting with the BLM-DNA2 complex. To further consolidate this finding, we sought to test whether GNL3 itself binds to ssDNA. We developed a PLA-based assay, referred to as ssDNA-PLA, to detect the interaction of GNL3 protein with BrdU-incorporated ssDNA. U2OS cells were first cultured in the presence of BrdU for 20 hours and treated with ETO or control DMSO for 2 hours. The cells were stained with anti-BrdU and anti-GNL3 antibodies followed by PLA assays under non-denaturing conditions, in which anti-BrdU only recognizes BrdU-incorporated ssDNA, but not double-stranded (ds) DNA. As shown in Figs. 4j and 4k, ETO treatment markedly induced the association of GNL3 with BrdU-incorporated ssDNA. Thus, GNL3 interacts with ssDNA upon DSBs, possibly generated by MRN-CtIP-mediated short-range DNA end resection.

**Figure 4.**
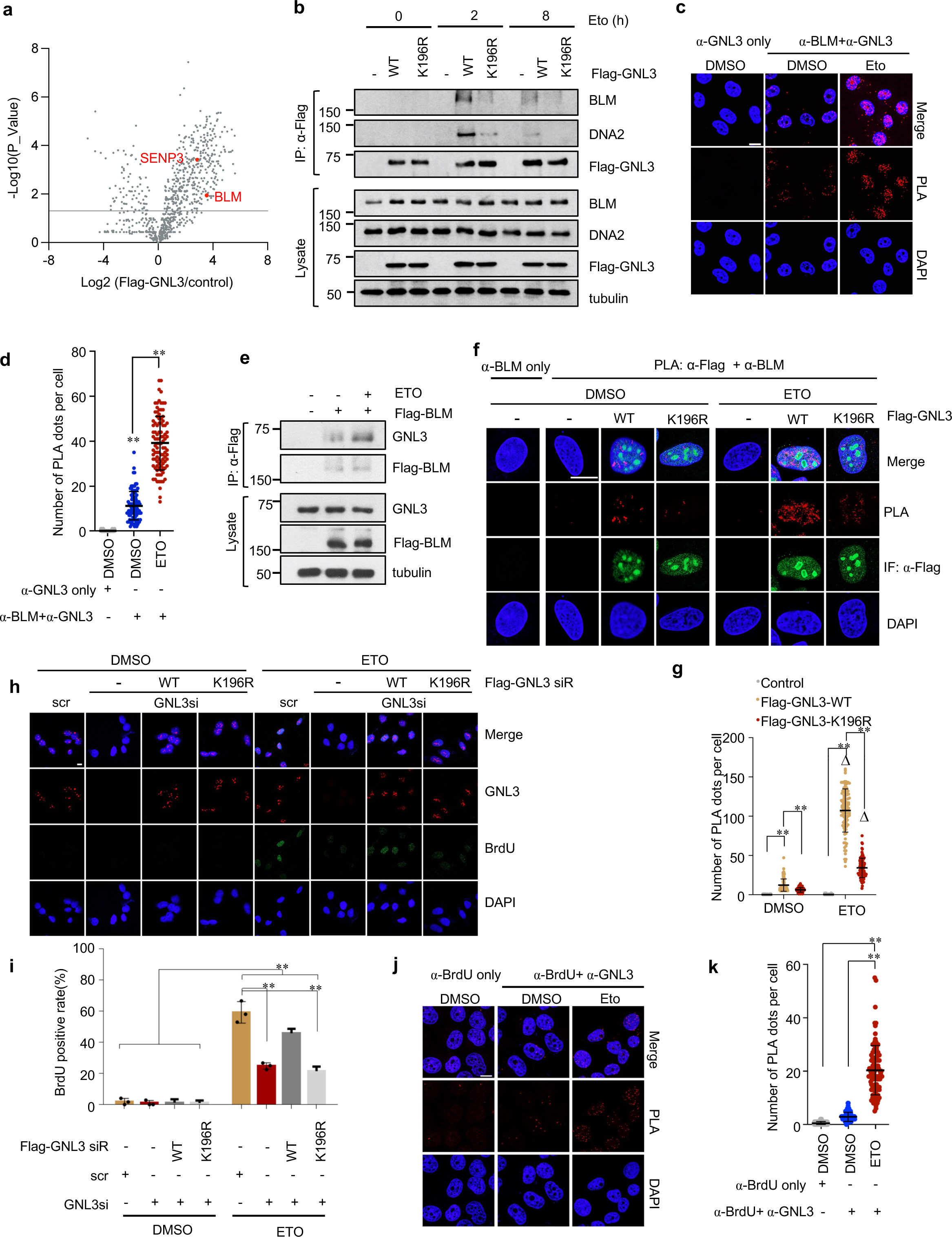
GNL3 regulates DNA end resection by interacting with BLM-DNA2. **(a)** Volcano plot of quantitative MS analysis of affinity-purified Flag-GNL3-associated protein complex from triplicate experiments. X-axis represents the Log2 ration of peptide counts from elutes of 293-Flag-GNL3 immunoprecipitates to those from 293 cells expressing a vector control. The significant cut line indicated is set at P<0.05. (**b)** GNL3 interacts with BLM-DNA2 in response to DNA damage dependently on its SUMOylation. U2OS cells transfected with the indicated plasmids were treated with 20 μM ETO for the indicated times and assayed by co-IP using anti-Flag antibody followed by IB. (**c)(d)** Interaction between endogenous GNL3 and BLM is also increased upon DNA damage. U2OS cells treated with control or 20 μM ETO for 2 hours were assayed by PLA using anti-GNL3 and anti-BLM antibodies. Anti-GNL3 only staining is used as a control. Shown are representative confocal images (c) and the quantification (d). **(e)** U2OS cells transfected with Flag-BLM and treated with control or 20 μM ETO for 2 hours were assayed by co-IP using anti-Flag antibody followed by IB. **(f)(g)** GNL3 SUMOylation at K196 is required for interaction with BLM upon DNA damage. U2OS cells transfected with Flag-GNL3 WT or the K196R mutant were treated with control or 20 μM ETO for 2 hours and assayed by PLA using anti-Flag and anti-BLM antibodies. Anti-BLM only staining is used as a control. Shown are representative confocal images (f) and the quantification (g). **(h)(i)** GNL3 SUMOylation regulates DNA end resection in response to DNA damage. U2OS cells transfected with scr or GNL3 siRNA together with Flag-tagged siRNA-resistant WT GNL3 or the K196R mutant were cultured in the presence of 10 μM BrdU for 24 hours, followed by IF staining with anti-BrdU and anti-GNL3 under native conditions. Shown are representative IF images (h) and the quantification (i). **(j)(k)** GNL3 interacts with ssDNAs. U2OS cells cultured in the presence of 10 μM BrdU for 24 hours were treated with control or 20 μM ETO for 2 hours and assayed by PLA using anti-BrdU and anti-GNL3 antibodies under native conditions. Shown are representative IF images (j) and the quantification (k). **P<0.01 compared to controls. △P<0.01, compared to the respective DMSO-treated group (triangle symbol). Scale bar, 10 μm.

### SUMO-SIM interaction directs the binding between GNL3 and BLM

To understand how GNL3 SUMOylation regulates its interaction with BLM, we reasoned that the interaction involves SUMO-SUMO-interacting motif (SIM) interactions. SIMs are characterized by a short stretch of core hydrophobic amino acids (V/I)x(V/I)(V/I) flanked by acidic residues^76^. It has been shown that BLM SUMOylation at K317 and K331 is necessary for its interaction with RAD51 and HR repair^32,33^. BLM also contains two SIMs, ^217^QIDL^2^^20^ and ^235^VICI^2^^38^, which are required for its SUMO binding and SUMOylation^77^. Similarly, RAD51 can be SUMOylated at K57 and K70 by TOPORS^38^ and contains a SIM, ^261^VAVV^2^^64^. Both RAD51 SUMOylation and its SIM are required for its function in HR repair^38,39^. RAD51 binds to SUMO and SUMOylated BLM^33^. Thus, BLM SUMOylation plays a critical role in DNA end resection and RAD51 loading by SUMO- and SIM-mediated BLM-RAD51 interaction. Since SUMOylation of GNL3 promotes its interaction with both BLM and RAD51, it is conceivable that GNL3 SUMOylation regulates its interaction with BLM and RAD51 via SUMO-SIM interactions. To test whether GNL3 also contains SIMs, we mutated a putative SIM predicted by silico analysis, ^143^VVLEVL^1^^49^ to ^143^V<u>AA</u>E<u>A</u>L^1^^49^ (Fig. 5a) and found that the GNL3^SIMm^ mutant failed to interact with either endogenous SUMO2 (Fig. 5b and supplementary Fig. 5a) or ectopically expressed non-conjugatable form of SUMO2, SUMO2G, that contains only one Gly residue at the C-terminus (Fig. 5c), and abolished GNL3 SUMOylation (Fig. 5d), confirming that the sequence is an authentic SIM. Notably, the GNL3^SIMm^ mutant failed to rescue γH2AX foci formation upon DNA damage induced by knockdown of endogenous GNL3 (Figs. 5e & 5f) and significantly attenuated the GNL3 interaction with BLM and DNA2 (Figs. 5g-5i & Supplementary Fig. 5b). These results suggest that the GNL3 SIM is critical for mediating interaction with BLM and function in HR via SUMO-SIM interactions. To further test this possibility, we generated a BLM mutant with mutations of both of its SIMs, BLM^SIMdm^, and found that mutating the two SIMs in BLM markedly reduced its interaction with GNL3 in response to ETO treatment as determined by both co-IP (Fig. 5j) and PLA (Figs. 5k, 5l & Supplementary Fig. 5c) assays. We also generated the SUMOylation-defective K317R and K331R BLM mutant, BLM^2KR^, and observed a marked attenuation of the binding of GNL3 to the BLM^2KR^ mutant compared to WT BLM in response to ETO treatment as determined by both co-IP (Fig. 5m) and PLA (Figs. 5n, 5o & Supplementary Fig. 5d) assays. Similar results were observed using GFP-BLM^2KR^ mutants (Supplementary Fig. 5e). Together, these results suggest that the GNL3-BLM association involves two SUMO-SIM interactions (Fig. 5p).

**Figure 5.**
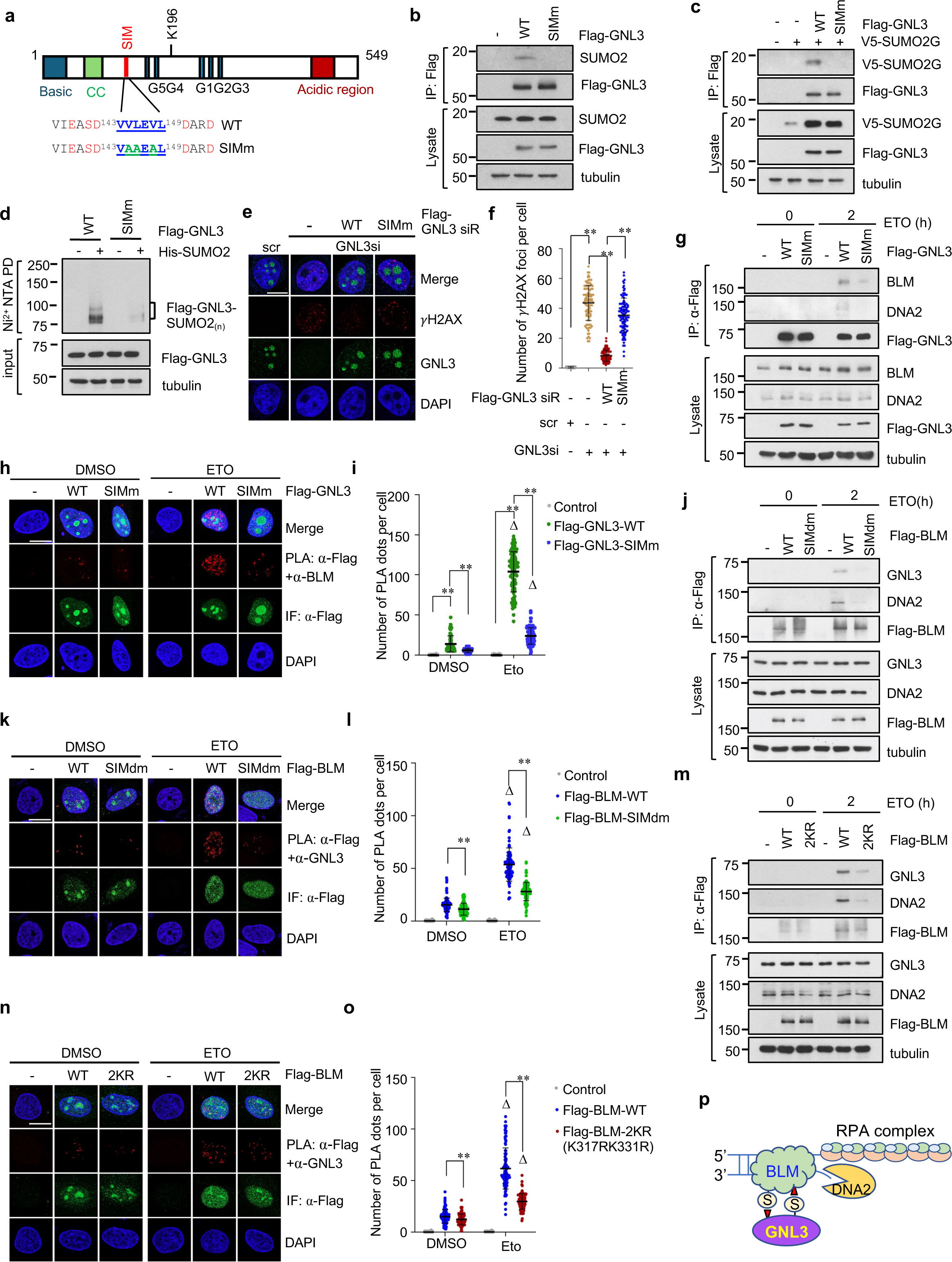
SUMO-SIM interaction directs the GNL3 interaction with BLM. **(a)** A diagram of GNL3 showing the sequences of the putative SIM sequence and the mutated SIM. Underlined are the SIMs flanked by acidic residues. **(b) (c)** Mutating the SIM abolishes GNL3 interaction with SUMO2. U2OS cells transfected with WT GNL3 or the SIM mutant (b) or together with V5-SUMO2G (c) were assayed by co-IP using anti-Flag antibody, followed by IB. **(d)** Mutating the SIM attenuates GNL3 SUMOylation. H1299 cells transfected with the indicated plasmids were subjected to Ni^2+^-NTA PD to detect SUMOylated GNL3. **(e)(f)** Ectopic expression of WT GNL3, but not the SIM mutant, abolishes γH2AX foci formation upon knockdown of endogenous GNL3. U2OS cells transfected with scr or GNL3 siRNA together with Flag-tagged siRNA-resistant WT GNL3 or the SIMm mutant were assayed by IF staining with anti-GNL3 and anti-γH2AX. Shown are representative confocal images (e) and the quantification (f). **(g)-(i)** Mutating the SIM attenuates GNL3 interaction with BLM in response to DAN damage. U2OS cells transfected with Flag-GNL3 WT or the SIM mutant were treated with DMSO or 20 μM ETO for 2 hours and assayed by co-IP using anti-Flag antibody (g) and PLA using anti-Flag and anti-BLM antibodies (h)(i). Shown are representative confocal images (h) and the quantification (i). **(j)-(l)** BLM SIM is critical for its interaction with GNL3 in response to DNA damage. U2OS cells transfected with Flag-BLM WT or the mutant with mutations of both SIMs (SIMdm) were treated with control or 20 μM ETO for 2 hours and assayed by co-IP using anti-Flag antibody (j) and PLA using anti-Flag and anti-BLM antibodies (K)(L). Shown are representative confocal images (k) and the quantification (l). **(m)-(o)** BLM SUMOylation is also critical for its interaction with GNL3 in response to DNA damage. U2OS cells transfected with Flag-BLM WT or the K317R;K331R (2KR) mutant were treated with control or 20 μM ETO for 2 hours and assayed by co-IP using anti-Flag antibody (m) and PLA using anti-Flag and anti-GNL3 antibodies (n)(o). Shown are representative confocal images (n) and the quantification (o). **(p)** Schematic diagram showing the interaction of GNL3 and BLM via their SUMO-SIM interactions. Scale bar, 10 μm. **P<0.01 compared to controls. △P<0.01, compared to the respective DMSO-treated group (triangle symbol).

### USP36 is a SUMO ligase for GNL3

To identify SUMO ligases that mediate GNL3 SUMOylation, we examined the PIAS family SUMO ligases and the nucleolar SUMO ligase USP36, a nucleolar deubiquitinase also acting as a SUMO ligase discovered by us^66–69^. We found that USP36 markedly promoted GNL3 SUMOylation, while PIAS1, but not other PIAS family members, slightly promoted GNL3 SUMOylation (Fig. 6a and Supplementary Fig. 6a), suggesting that USP36 is a major GNL3 SUMO ligase. Consistently, GNL3 was identified in our previously purified USP36 complex^67^ (Fig. 6b). We confirmed the USP36-GNL3 interaction by co-IP (Figs. 6c & 6d). PLA showed that the endogenous USP36-GNL3 interaction is significantly increased upon ETO treatment (Figs. 6e & 6f). RNase treatment did not disrupt the interaction between USP36 and GNL3, whereas USP36 interaction with ribosomal protein L30 (RPL30) was largely abolished (Supplementary Fig. 6b), suggesting that USP36 interacts with GNL3 independently of RNA and intact ribosome. Using co-IP assays, we further mapped that both the N-terminal USP domain and the C-terminal domain of USP36 are capable of binding to GNL3 (Supplementary Fig. 6c & 6d). Immunofluorescence (IF) staining (Supplementary Fig. 6e) and cell fractionation assays (Supplementary Fig. 6f) showed that both USP36 and GNL3 were found to predominantly localize in the nucleolus. In supporting its role in SUMOylating GNL3, knockdown of USP36 significantly reduced GNL3 interaction with the BLM-DNA2 complexes (Fig. 6g). Thus, USP36 is a likely novel regulator of GNL3-mediated DDR pathway.

**Figure 6.**
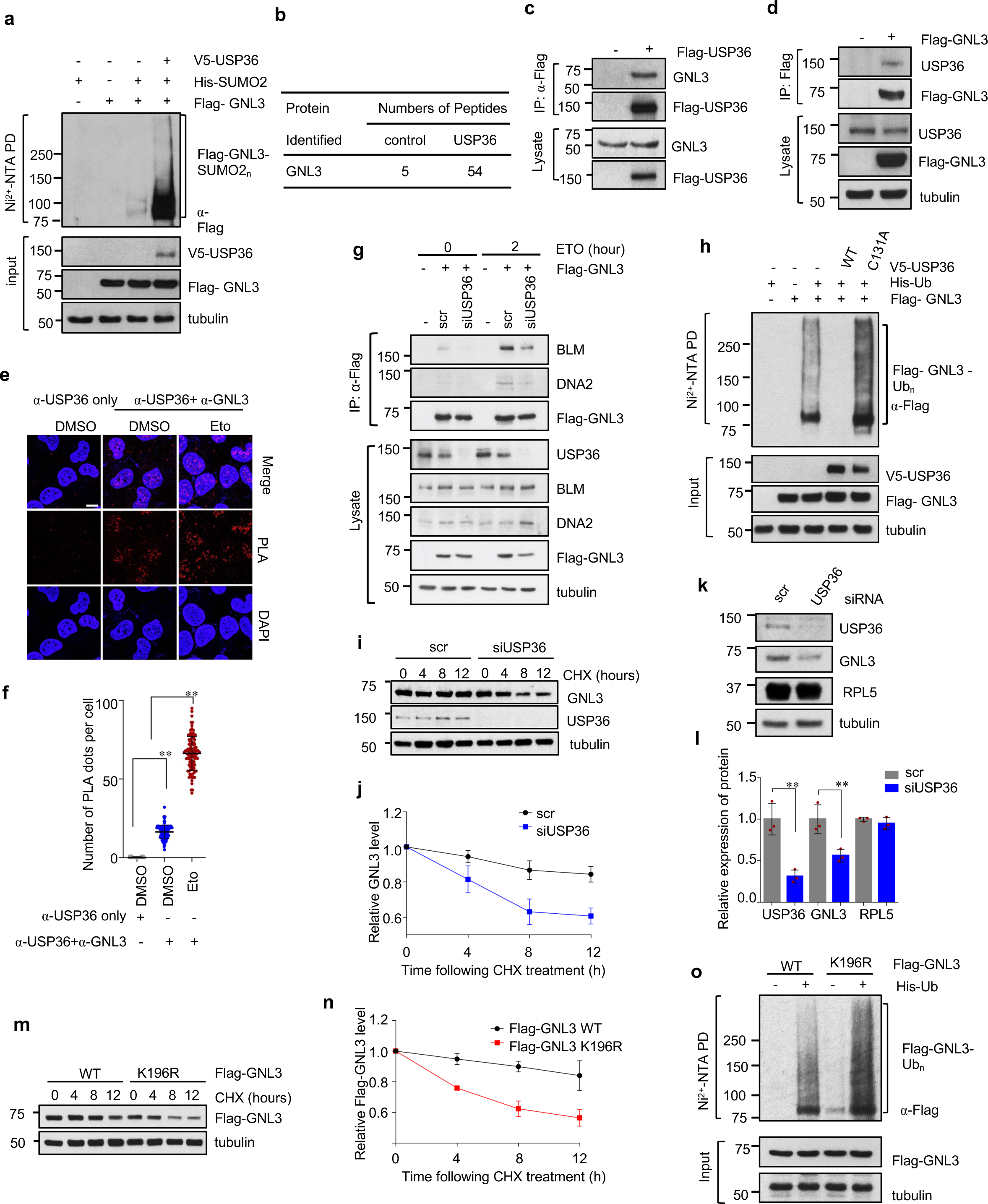
USP36 is a GNL3 SUMO ligase. **(a)**. USP36 promotes GNL3 SUMOylation in cells. H1299 cells transfected with the indicated plasmids were assayed by Ni^2+^-NTA PD under denaturing conditions followed by IB. The SUMO2-modified GNL3 is indicated. The protein expression is shown in the bottom panels. **(b)**. GNL3 peptide identified in affinity-purified Flag-USP36 complex compared to control from 293 cells. **(c)(d)** Co-IP of Flag-USP36 and GNL3 in cells. 293 cells transfected with Flag-USP36 (c) or Flag-GNL3 (d) were assayed by co-IP using anti-Flag antibody. **(e)(f)**. DNA damage enhances USP36-GNL3 interaction. U2OS cells treated with DMSO or 20 μM ETO for 2 hours were assayed by PLA using anti-USP36 and anti-GNL3 to detect the interaction of endogenous USP36 and GNL3. Shown are representative IF images (e) and the quantification (f). Scale bar, 10 μm. **P<0.01**. (g)** Knockdown of USP36 attenuates GNL3 interaction with BLM-DNA2 in response to DAN damage. U2OS cells transfected with Flag-GNL3 together with scrambled control or USP36 siRNA were treated with DMSO or 20 μM ETO for 2 hours and assayed by co-IP using anti-Flag antibody. (**h**) USP36 deubiquitinates GNL3. H1299 cells transfected with the indicated plasmids were assayed by Ni^2+^-NTA-PD to detect the ubiquitinated GNL3. **(i)(j**) USP36 regulates GNL3 protein stability. U2OS cells transfected with scr or USP36 siRNA were treated with cycloheximide (CHX) for different times and assayed by IB (i). Quantification is shown in (j). **(k)(l**) Knockdown of USP36 reduces GNL3 levels. U2OS cells transfected with scr or USP36 siRNA were assayed by IB (k). Quantification is shown in (l). **P<0.01, compared to scr controls. **(m)(n**) Mutating K196 reduces GNL3 stability. U2OS cells transfected with WT GNL3 or the K196R mutant were treated with CHX for different times and assayed by IB (m). Quantification is shown in (n). (**o**) Mutating K196 increases GNL3 ubiquitination. H1299 cells transfected with the indicated plasmids were assayed by Ni^2+^-NTA PD to detect the ubiquitinated GNL3.

### USP36 deubiquitinates GNL3 and regulates its levels

As USP36 is a deubiquitinating enzyme (DUB), we next examined whether it can deubiquitinate GNL3. We transfected cells with Flag-GNL3 along with either WT USP36 or its catalytically inactive mutant (C131A). In vivo ubiquitination assays under denaturing conditions showed that WT USP36, but not the C131A mutant, completely abolished GNL3 ubiquitination (Fig. 6h).

Consistently, knockdown of USP36 significantly decreased the stability of GNL3 as determined by half-life assays (Figs. 6i & 6j) and reduced the levels of GNL3, but not RPL5 (Figs. 6k & 6l). Thus, USP36 acts as a SUMO ligase/DUB dual-function enzyme for GNL3. Interestingly, the SUMOylation-defective K196R mutant showed reduced protein stability (Figs. 6m & 6n) and increased ubiquitination (Fig. 6o), suggesting that K196 SUMOylation inhibits GNL3 ubiquitination and proteasome-mediated degradation.

### SENP3 deSUMOylates GNL3 and is also required for DNA damage repair

DeSUMOylation of DDR proteins by SUMO proteases such as SENP3 and SENP6 has been shown to regulate DSB repair^30,34^. Given that both SENP3 and GNL3 are predominantly localized to the nucleolus and shuttle between the nucleus and the nucleolus^51,78–80^, and that SENP3 was also present in our purified Flag-GNL3-associated protein complexes (Fig. 4a), we examined whether SENP3 could regulate GNL3. We confirmed that SENP3 interacts with GNL3 by co-IP (Figs. 7a & 7b). The interaction between SENP3 and GNL3 is also RNA-independent (supplementary Figs. 7a & 7b). The N-terminal domain of SENP3 is capable of binding to GNL3 (supplementary Figs. 7c & 7d). Of note, SENP3 and GNL3 co-localized mainly in the nucleolus that was not affected by the ETO treatment (Fig. 7c). PLA assays further confirmed the endogenous interaction between GNL3 and SENP3 (Fig. 7d & 7e). Next, we examined whether SENP3 deSUMOylates GNL3. As shown in Fig. 7f, WT SENP3, but not its catalytically inactive C532S mutant, markedly deSUMOylated GNL3. Thus, SENP3 is a SUMO protease for GNL3. SENP3 knockdown did not affect the nucleolar localization of either WT GNL3 or the K196R mutant (Supplementary Fig. 7e). As SUMOylation increases GNL3 stability (Figs. 6i-j, 6m-n), we examined whether SENP3 deSUMOylation would destabilize GNL3. However, knockdown of SENP3 did not significantly affect the total levels of GNL3 (Supplementary Figs. 7f & 7g), suggesting that SENP3 does not impact the total levels and localization of GNL3, although we cannot exclude the possibility of a small fraction of GNL3 in chromatin may be subjected to SUMOylation-regulated turnover.

**Figure 7.**
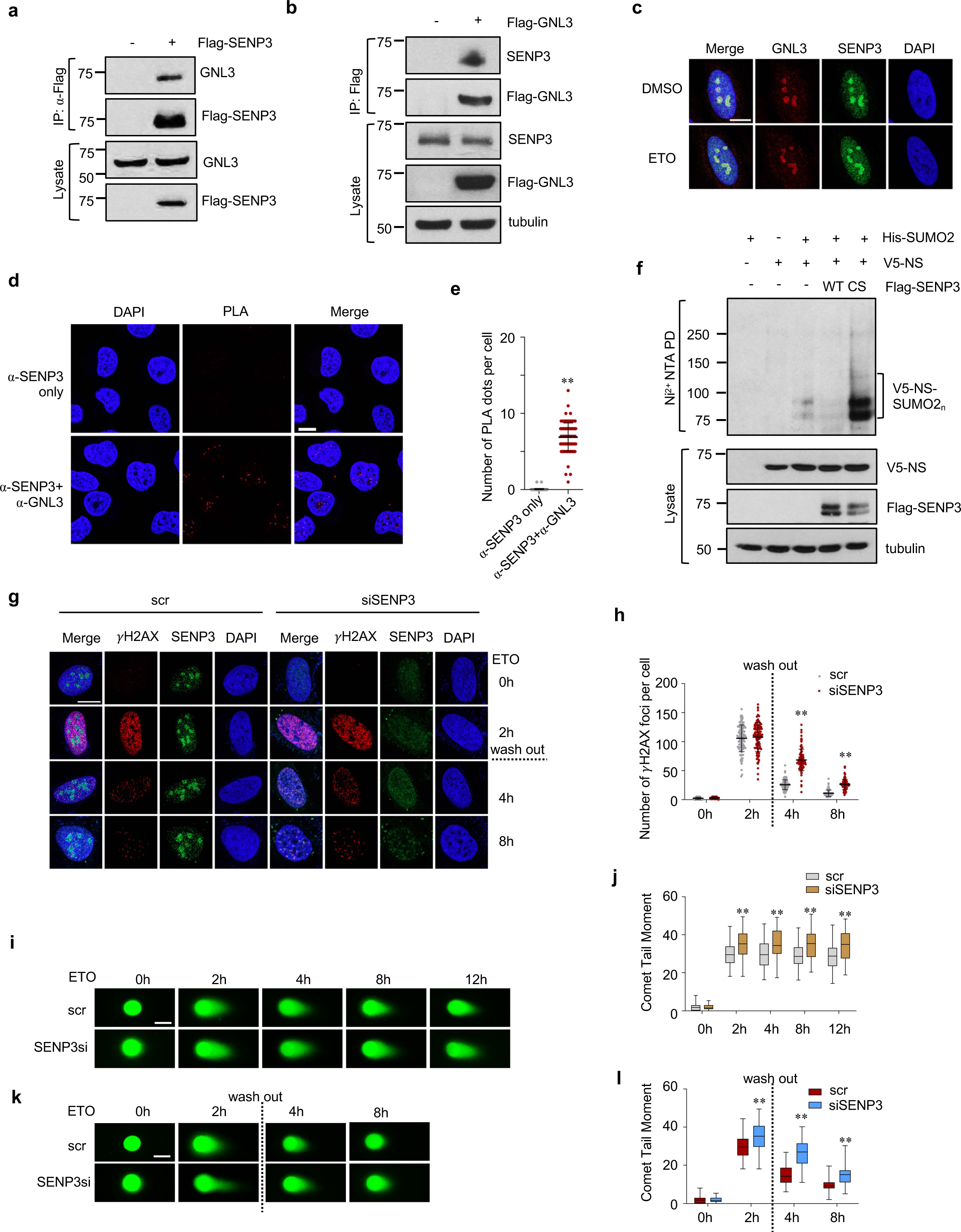
SENP3 deSUMOylates GNL3. **(a)(b)** Co-IP of SENP3 and GNL3 in cells. 293 cells transfected with Flag-SENP3(a) or Flag-GNL3 (b) were assayed by co-IP using anti-Flag antibody. (**c**) SENP3 co-localizes with GNL3 in the nucleolus. U2OS cells were treated with DMSO or 20 μM ETO for 2 hours and assayed by IF staining with anti-SENP3 and anti-GNL3 antibodies. (**d**)(**e**) Endogenous SENP3 interacts with GNL3 determined by PLA assays using anti-SENP3 and anti-GNL3 antibodies. Anti-SENP3 only is used as a control. Shown are representative PLA images (d) and the quantification (e). (**f**) SENP deSUMOylates GNL3. H1299 cells transfected with the indicated plasmids were subjected to Ni^2+^-NTA PD followed by IB to detect SUMOylated GNL3. **(g)(h)**. Knockdown of SENP3 attenuates the disappearance of γH2AX foci upon DNA damage. U2OS cells transfected with scr or SENP3 siRNA were treated without or with 20 μM ETO for 2 hours followed by washing out and replacing with fresh medium and fixed at the indicated time. The cells were stained with anti-γH2AX and anti-SENP3. Shown are representative IF images (g) and the quantification of the γH2AX foci (h). **(i)-(l)**. Knockdown of SENP3 attenuates DNA damage repair. Comet assays were conducted in U2OS cells transfected with scr or SENP3 siRNA were treated without or with 20 μM ETO for different times indicated (i)(j) or the medium was washed out and replaced with fresh medium and harvested at the indicated time (k) (l). Shown are the representative images (i) (k) and quantification (j)(l) of Comet tailor moment. Scale bar, 10 μm. **P<0.01, compared to controls.

We next sought to determine whether SENP3 regulates DNA damage repair. As shown in Figs. 7g and 7h, knockdown of SENP3 markedly attenuated the clearance of γH2AX foci upon ETO treatment, indicative of an impaired DNA damage repair. Comet assays showed that knockdown of SENP3 resulted in increased tail moment compared to the scrambled control, both during ETO treatment and after washing out (Fig. 7i–7l), suggesting that SENP3 knockdown reduces the efficiency of DNA damage repair. Repair reporter assays also showed that knockdown of SENP3 reduced both HR and NHEJ repair activities (Supplementary Fig. 7h). These results indicate that SENP3 is essential for DNA damage repair, consistent with the recently reported function of SENP3 in DNA damage repair by deSUMOylating TIP60, Mre11 and NPM^30,81–83^.

### GNL3 implication in breast cancer

To examine whether the role of GNL3 and its SUMOylation in HR repair is implicated in tumorigenesis and therapeutic response, we focused on breast cancer, as TCGA data showed that GNL3 mRNA expression is significantly higher in primary breast cancers compared to normal control (Fig. 8a) and significantly higher in triple-negative breast cancers (TNBCs) compared to non-TNBCs (Fig. 8b). Data from NCI Clinical Proteomic Tumor Analysis Consortium (CPTAC) database showed that GNL3 protein is significantly higher in breast cancers compared to normal control (Fig. 8c). High GNL3 protein expression is associated with poor patient survival in two available public datasets (Figs. 8d & 8e). Consistent with the role of GNL3 in HR repair, its high expression is significantly associated with DNA repair hallmark (Fig. 8f & Supplementary Fig. 8a). Collectively, these data suggest that GNL3 is overexpressed in BCs and its role in HR could contribute to therapeutic resistance. Thus, we next examined whether GNL3 SUMOylation plays a role in breast cancer therapeutic response. Consistent with the critical role for GNL3 and its SUMOylation in HR repair, knockdown of GNL3 also induced the γH2AX foci formation (Fig. 8g) and levels (Figs. 8h & 2f) in TNBC MDA-MB-231 cells, which can be abolished by expression of WT GNL3, but not the K196R mutant (see Fig. 2g).

**Figure 8.**
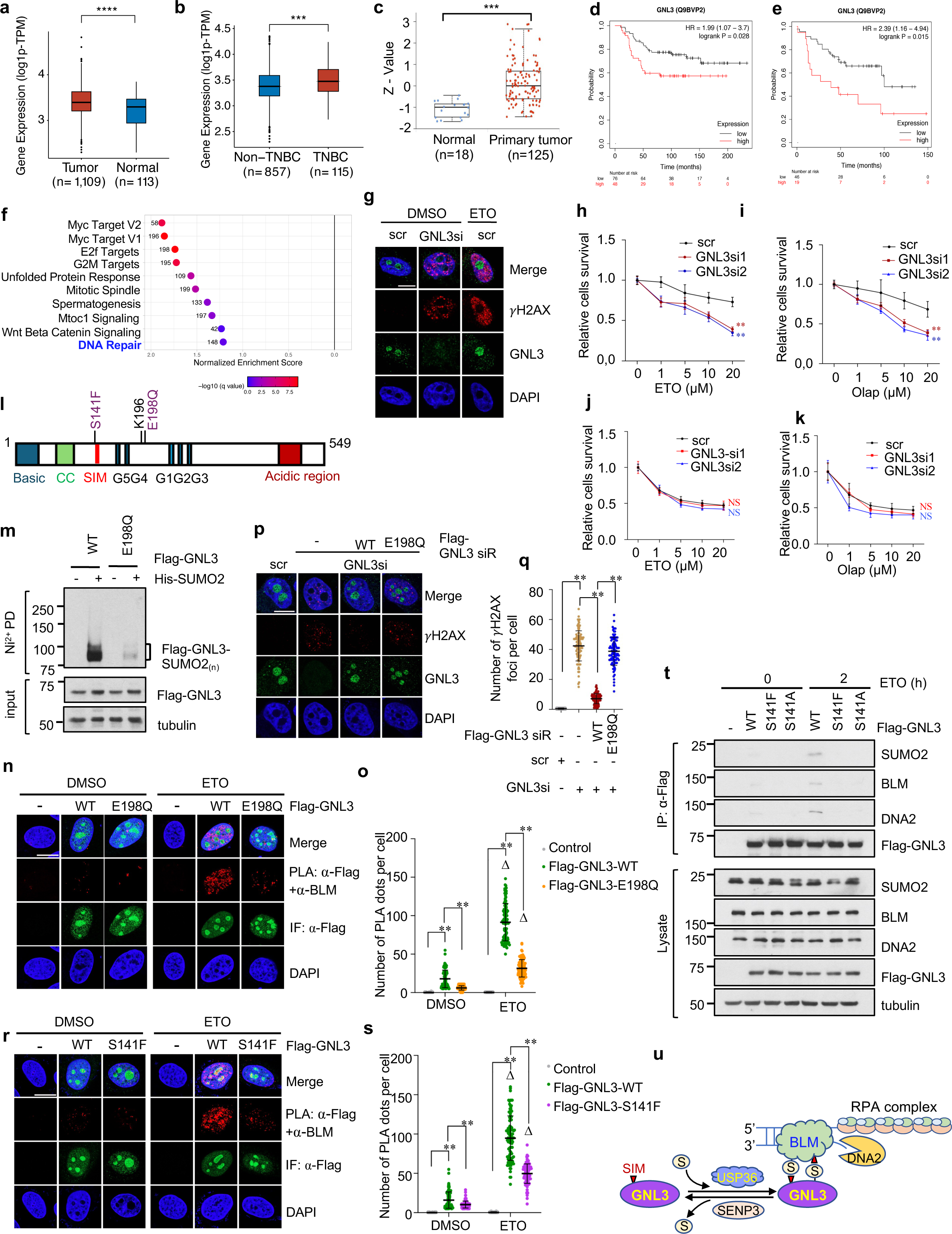
GNL3 is implicated in breast cancer. **(a)(b)** GNL3 mRNA expression is significantly increased in BC compared to normal controls (a) and increased in TNBC compared to non-TNBC (b) (data from TCGA). ***P<0.001; ****P<0.0001. **(c)** GNL3 protein expression is increased in BC compared to normal controls (data from the NCI Clinical Proteomic Tumor Analysis Consortium (CPTAC) database). (**d**)**(e)**. High expression of GNL3 mRNA (d) and protein (e) is associated with poor patient survival (data from Kaplan-Meier plotter). **(f)** GNL3 expression is associated with the DNA damage hallmark in TNBC. Shown are the top 10 hallmarks from TCGA data. **(g)** Knockdown of GNL3 induces DNA damage in breast cancer cells. MDA-MB-231 cells were transfected with scr or GNL3 siRNA or treated with 20 μM ETO for 2 hours and stained with anti-γH2AX for detecting γH2AX foci (g). **(h)-(k).** Knockdown of GNL3 sensitizes HR-proficient, but not HR-deficient, breast cancer cells to treatment with DNA damage agents. MDA-MB-231 (h & i) and MDA-MB-436 (j & k) cells transfected with scr or GNL3 siRNAs were treated with different doses of ETO (h & j) or Olaparib (i & k), followed by MTT assays. *P<0.01 compared to scr control. **(l)** Diagram of GNL3 showing the breast cancer-derived mutations E198Q and S141F. **(m)** The E198Q mutant attenuates GNL3 SUMOylation. H1299 cells transfected with the indicated plasmids were subjected to Ni^2+^-NTA PD to detect SUMOylated GNL3. **(n)(o)** The E198Q mutant GNL3 showed attenuated interaction with BLM in response to DNA damage. U2OS cells transfected with Flag-GNL3 WT or the E198Q mutant were treated with control or 20 μM ETO for 2 hours and assayed by PLA using anti-Flag and anti-BLM antibodies. Shown are representative confocal images (n) and the quantification of PLA signals (o). **(p)(q)** Ectopic expression of WT GNL3, but not the E198Q mutant, abolishes γH2AX foci formation upon knockdown of endogenous GNL3. U2OS cells transfected with scrambled (scr) or GNL3 siRNA together with Flag-tagged siRNA-resistant (siR) WT GNL3 or the E198Q mutant were assayed by IF staining with anti-GNL3 and anti-γH2AX. Shown are representative confocal images (p) and the quantification (q) of the γH2AX foci. **(r)(s)** The S141F mutant GNL3 showed attenuated interaction with BLM in response to DNA damage. U2OS cells transfected with Flag-GNL3 WT or the S141F mutant were treated with control or 20 μM ETO for 2 hours and assayed by PLA using anti-Flag and anti-BLM antibodies. Shown are representative confocal images (r) and the quantification of PLA signals (s). **(t)** Mutating S141 attenuates GNL3 interaction with BLM-DNA2 and SUMO2 in response to DNA damage. U2OS cells transfected with Flag-GNL3 WT or the S141F or S141A mutant were treated with control or 20 μM ETO for 2 hours and assayed by co-IP to detect the GNL3 interaction with BLM-DNA2 and SUMO2. **(u)** Schematic diagram of the working model showing that USP36 and SENP3-regulated GNL3 SUMOylation regulates GNL3 interaction with BLM via their SUMO-SIM interactions. Scale bar, 10 μm. **P<0.01 compared to controls. △P<0.01, compared to the respective DMSO-treated group (triangle symbol).

Consistently, knockdown of GNL3 significantly increased the sensitivity of HR competent MDA-MB-231 (Figs. 8h & 8i), but not the HR deficient MDA-MB-436 (Figs. 8j, 8k & Supplementary Fig. 8b), cells to ETO and Olaparib, suggesting the GNL3 could be a promising therapeutic target in HR-proficient cancers.

Interestingly, several GNL3 variants with unknown significance (VUS) in the SUMO and SIM motifs are present in TCGA breast cancer patients, including E198Q (abolishing SUMO site sequence consensus) and S141F (reducing negative charge and likely weakening the SIM) (Fig. 8l). Mutating E198 to Gln (Q) abolished GNL3 SUMOylation (Fig. 8m) and attenuated GNL3 interaction with BLM upon ETO treatment as determined by PLA (Figs. 8n, 8o & Supplementary Fig. 8c). The E198Q mutant fails to rescue DNA damage response induced by knockdown of endogenous GNL3 (Figs. 8p & 8q), suggesting the E198Q mutation inactivates the GNL3 SUMOylation pathway and fails to function in HR repair. Additionally, mutating S141 also markedly reduced GNL3 binding to BLM and SUMO2 (Figs. 8r-8t & Supplementary Fig. 8d), suggesting that mutating S141 impairs the SIM function and GNL3 function in HR. Together, our results suggest that these VUSs attenuate the SUMO regulation of GNL3 and HR repair and may contribute to mammary tumorigenesis.

## Discussion

Long-range DNA end resection by the BLM-DNA2 helicase-nuclease complex is crucial for DSB HR repair. However, how this critical enzymatic process is regulated in cells is not well understood. In this study, we discovered that GNL3 plays a critical role in HR repair via interacting with BLM-DNA2 and regulating DNA end resection. Mechanistically, this process is tightly controlled by SUMOylation. We found that GNL3 is SUMOylated at K196 in response to DSBs and it contains a SIM. Both SUMOylation and the SIM are required for GNL3 to bind to BLM and function in HR. Notably, BLM contains two SIMs and is also regulated by SUMOylation ^32,33,77^. We showed that either abolishing BLM SUMOylation or mutating the two SIMs impairs its interaction with GNL3. Together, these data demonstrate that the GNL3-BLM interaction is mediated by mutual SUMO-SIM interactions (Fig. 8v). In response to DSBs, GNL3 SUMOylation is markedly increased, which promotes its interaction with the BLM-DNA2 complex. Thus, GNL3 acts as a critical regulator of the HR pathway by fine-tuning DNA end resection mediated by the BLM-DNA2 complex.

Supporting its role in regulating DNA end resection, GNL3 localizes to sites of DNA damage, as indicated by its association with γH2AX and ssDNA. The GNL3-ssDNA intermediates may play a critical role in regulating BLM-DNA2 recruitment upon DSBs by stabilizing the short-range ssDNAs generated by the MRN-CtIP. Through its interaction with BLM, GNL3 may also directly stimulate BLM helicase activity. Both RPA ^84^ and CtIP^85^ have been shown to interact with BLM and enhance its helicase activity. Chromatin-associated factors such as recently reported ZNF280A promote long-range DNA-end resection by facilitating the recruitment of the BLM-DNA2 complex to DNA DSB sites and enhancing its activity ^86^. The BLM complex includes additional components such as BMI1, BMI2, and TOP3A ^87^. Interestingly, both BMI1 and TOP3A have been shown to be regulated by SUMOylation^88–91^. Thus, it will be interesting to examine whether SUMOylation regulates GNL3 interaction with other BLM complex components, BMI1, BMI2, and TOP3A, and whether DNA damage triggers the coordinated BLM complex-GNL3 protein group SUMOylation. In addition, BLM plays a critical role in RAD51 loading through SUMO- and SIM-mediated BLM-RAD51 interaction^31^ and it promotes synapsis during HR by stabilizing RAD51 in the displacement loop (D-loop) upon template strand invasion ^92^. We demonstrated that both GNL3-RAD51 interaction and RAD51 foci formation following DSBs also depend on GNL3 SUMOylation, suggesting that GNL3- RAD51 interaction may play a direct role in RAD51 loading and stabilization within the D-loop. Future studies are warranted to elucidate the precise mechanism by which GNL3-BLM interaction contributes to DNA end resection and HR repair.

Intriguingly, we found that several breast cancer-derived VUSs, including E198Q and S141F, impair the SUMOylation-regulated GNL3-BLM interaction. This suggests that these variants could contribute to mammary tumorigenesis by impairing HR repair and promoting genomic instability, thus being pathogenic mutations in breast cancer. This finding further reinforces the critical role of GNL3-SUMO regulation in HR repair. Notably, depletion of GNL3 sensitizes HR proficient MDA-MB-231, but not HR-deficient MDA-MB-436, cells to treatment with ETO and Olaparib, suggesting that GNL3 could be a promising therapeutic target in HR proficient and GNL3 overexpressing cancers. As GNL3 is an essential gene required for animal development and stem cell maintenance^51,93,94^, targeting its SUMOylation may represent a promising approach to sensitize cancer cells to DNA-damaging agents while potentially minimizing effects on normal cellular functions. In this context, it is imperative to determine whether GNL3 SUMOylation is also essential for its function in cell growth and cell cycle regulation, particularly through its regulation of ribosome biogenesis or the cell cycle.

GNL3 is a nucleolar-nucleoplasm shuttling protein that predominantly localizes to the nucleolus. DNA damage or abolishing GNL3 SUMOylation does not apparently alter its predominant nucleolar localization. GNL3 has been shown to interact with γH2AX in the nucleoplasm of S-phase cells ^63^. Our PLA assays revealed that the GNL3-BLM interactions occur throughout the nucleus. These observations suggest that a subset of SUMOylated GNL3 in the nucleoplasm may be critical for HR. Additionally, nucleolar integrity is known to be impaired by most, if not all, cellular stressors, including DNA damage ^95^. Thus, subtle changes in nucleolar dynamics may be sufficient for GNL3 to interact with BLM and regulate DSB repair via HR, especially during S and G2 cell cycle phases when the HR pathway functions. Yet, this does not exclude the possibility that the nucleolar localization of GNL3 may be critical for HR, given that many DDR proteins transiently localize to the nucleolus ^96^. Particularly, we found that GNL3 SUMOylation is tightly regulated by the nucleolar SUMO ligase USP36 and SUMO protease SENP3 (see below).

USP36 SUMOylates several nucleolar ribosome biogenesis factors including the snoRNP complex components, EXOSC10 and Las1L, and plays a critical role in pre-rRNA processing and translation ^66,67,69^. We showed that USP36 also acts as a SUMO ligase for GNL3. Knockdown of USP36 significantly reduced GNL3-BLM interaction upon DSBs, suggesting that USP36 plays a role in DDR, at least in part by regulating GNL3. In addition, we showed that USP36 can interact with GNL3 in the nucleolus. Thus, USP36 may SUMOylate GNL3 in the nucleolus, thereby facilitating its interaction with BLM complex and promoting its shuttling to nuclear chromatin to execute HR function upon DSBs. Interestingly, USP36 also deubiquitinates GNL3 and regulates its protein levels, suggesting a critical crosstalk between SUMOylation and ubiquitination in regulating GNL3 levels. Thus, USP36 acts as DUB and SUMO ligase dual-function enzyme for GNL3 to regulate its SUMOylation and protein levels in response to DNA damage. GNL3 has recently been shown to play a role in overcoming DNA replication stress by sequestering DNA replication initiation factor ORC2 in the nucleolus, controlling origin firing, and thereby indirectly preventing nascent DNA resection by MRN-CtIP nucleases in response to replication stress ^65^. Given the role of GNL3 SUMOylation in regulating BLM-DNA2, our finding provides an alternative mechanism underlying a direct role for GNL3 in DNA damage repair in response to replication stress.

The role of SENP3 in DSB repair has been studied by several groups, particularly its function in deSUMOylating key DDR proteins including Mre11^30^, NPM^83^ and TIP60^81^. We showed that SENP3 also deSUMOylates GNL3, and that SENP3 depletion impairs DSB repair, as evidenced by the delayed γH2AX clearance. These findings suggest that SENP3-mediated deSUMOylation is essential for HR, potentially by facilitating the removal of the SUMOylated DDR proteins including GNL3 from the chromatin following repair. Since knockdown of SENP3 does not significantly affect total GNL3 protein levels, it remains to be tested whether SENP3 regulates the turnover of chromatin-associated GNL3 following the DNA damage via the ubiquitination-proteasome system. Future work would focus on understanding how USP36 and SENP3 regulate GNL3 SUMOylation in response to DNA damage and whether their activities are regulated by post-translational modifications such as ATM/ATR-mediated phosphorylation following DSBs.

In summary, we discovered that GNL3 SUMOylation is essential for protecting cells from spontaneous DNA damage as well as DSBs induced by exogenous agents. GNL3 has been shown to prevent telomere DNA damage by recruiting PML to SUMOylated telomeric repeat binding factor 1 (TRF1)^97^, raising the question of whether GNL3 SUMOylation is also required for this function. Together, our findings establish GNL3 as a novel regulator of DNA repair that safeguards genomic integrity through a SUMOylation-regulated mechanism.

## Methods

### Cell culture and reagents

Human U2OS, HEK293, HeLa, H1299, and MDA-MB-231 cells were cultured in Dulbecco’s modified Eagle’s medium (DMEM) supplemented with 10% (vol/vol) fetal bovine serum (FBS), 50 U/mL penicillin, and 0.1 mg/mL streptomycin at 37 °C in a 5% CO2 humidified atmosphere. These cell lines were obtained from ATCC and routinely monitored for mycoplasma contamination. Manufacturers performed authentication through short tandem repeat profiling. Etoposide (ETO), camptothecin (CPT), hydroxyurea (HU), cycloheximide (CHX), doxycycline (Dox), N-ethylmaleimide (NEM) and iodoacetamide (IAA) were purchased from Sigma-Aldrich. ML972 was purchased from Selleckchem. RNase A and RNaseT1 were purchased from Thermo Scientific.

### Plasmids, DNA transfections and site-directed mutagenesis

Flag-tagged GNL3 (Flag-GNL3) and Flag-BLM were cloned into pcDNA3-2Flag vector by PCR. The GNL3 cDNA was also subcloned into pcDNA3-V5 and pcDNA4-TO vectors to generate V5 tagged-GNL3 and tet-inducible pcDNA4-TO-2Flag-GNL3 plasmids, respectively. Flag-GNL3 siRNA resistant (Flag-GNL3-siR) plasmid was generated by site-directed mutagenesis using the QuikChange Kit (Agilent Technologies), in which the GNL3 siRNA targeting sequence (GNL3 si- 1) 5’-AAGAACTAAAACAGCAGCAGA-3’ was mutated to 5’-A<u>G</u>GA<u>GT</u>T<u>G</u>AA<u>G</u>CA<u>A</u>CA<u>A</u>CA<u>A</u>A-3’. The Flag-tagged GNL3 K196R, K275R, G261V mutants were generated by site-directed mutagenesis using Flag-GNL3 as a template. The Flag-tagged siRNA resistant GNL3 K196R, E198Q, S141F, S141A, and SIM mutants were generated using pcDNA4-TO-2Flag-GNL3-siR plasmid as a template. WT and siRNA-resistant GNL3 cDNAs were also cloned into pcDNA3 vector to generate untagged GNL3 plasmids. Flag-BLM with K317R;K331R mutant (Flag-BLM^2KR^) and with mutation of the two SIMs (BLM^SIMdm^) were also generated by site-directed mutagenesis. Flag-SENP3 and its deletion mutants were generated by PCR cloning. All the plasmids were confirmed by sequencing. V5-tagged SUMO2G plasmid with deletion of the last Gly residue was cloned by PCR. See Supplementary Table 1 for all other plasmids. Cells transfection was performed using TransIT®-LT1 reagents (Mirus Bio Corporation) or Lipofectamine 2000 (Life Technologies) reagents following the manufacturers’ protocol. Cells transfection was performed using TransIT®-LT1 reagents (Mirus Bio Corporation) or Lipofectamine 2000 (Life Technologies) reagents following the manufacturers’ protocol. Cells were harvested at 36-48 hours posttransfection for further analyses.

### Establishment of stable His10-SUMO2 and Flag-GNL3 expression cell lines

U2OS cells stably expressing His10-SUMO2 were generated by transient transfection with pcDNA3-His10-SUMO2 plasmid, followed by selection in medium containing 0.5 mg/ml neomycin (G418) for up to 2 weeks. Single colonies were isolated, expanded, and screened by immunoblot analysis using anti-His antibodies. 293 cells stably expressing control or Flag-GNL3 were established by transfecting with control pcDNA3-Flag and Flag-GNL3 plasmids, respectively, followed by selection in medium containing 0.5 mg/ml G418. Single colonies were also isolated, expanded, and screened by immunoblot analysis using anti-Flag antibodies. All the cells were maintained in cultured medium containing 0.5 mg/ml G418.

### Immunoblot (IB) and co-immunoprecipitation (co-IP), and antibodies

Cells were lysed in NP40 lysis buffer consisting of 50 mM Tris-HCl (pH 8.0), 0.5% Nonidet P-40, 1 mM EDTA, 150 mM NaCl, 1 mM phenylmethylsulfonyl fluoride (PMSF), 1 mM dithiothreitol (DTT), 1 μg/ml pepstatin A, and 1 mM leupeptin with brief sonication. Equal amounts of total protein were used for IB analysis. For Co-IP, cells were lysed in the above NP40 lysis buffer in the presence of 20 mM NEM and 5 mM IAA. Equal amounts of cell lysates were used for co-IP using anti-Flag (M2) antibodies agarose beads at 4°C for 4 hours. Beads were washed four times with lysis buffer and bound proteins were detected by IB using antibodies as indicated in the figure legends. A full list of antibodies and protocols can be found in Supplementary Table 2.

### Cell fractionation

Cell fractionation assays were performed essentially as described ^68^. Briefly, the freshly harvested cells were washed with PBS, resuspended in hypotonic buffer A (10 mM HEPES pH 7.8, 10 mM KCl, 1.5 mM MgCl_2_, 0.5 mM DTT) in the presence of PMSF and incubated for 10 minutes on ice. The cells were homogenized using a B pestle douncer followed by spinning down at 3,000 rpm for 5 minutes at 4 °C. The supernatant was supplemented with one-tenth volume of buffer B (0.3 M Tris-HCl pH 7.8, 1.4 M KCl, 30 mM MgCl_2_) and collected as the cytoplasmic fraction. The nuclear pellets were washed with buffer A once and then resuspended in buffer S1 (0.25 M sucrose, 10 mM MgCl_2_), layered over buffer S2 (0.35 M sucrose, 0.5 mM MgCl_2_), and centrifuged at 1,430 x g for 10 minutes at 4 °C. The pelleted nuclei were resuspended in buffer S2 with PMSF and sonicated using a microtip probe at a power setting of 50%. The sonicated nuclei were layered over buffer S3 containing 0.88 M sucrose and 0.5 mM MgCl_2_, and centrifuged at 3,000 x g for 10 min at 4 °C. The supernatant was collected as the nucleoplasm fraction. The pellet contained purified nucleoli and was dissolved in high salt RIPA buffer containing 50 mM Tris pH 7.5, 500 mM NaCl, 1% Nonidet P-40, 0.5% deoxycholate, and proteasome inhibitors in the presence of 80 U/mL DNase I on ice for 30 minutes. The lysates were then added with 2x volume of RIPA buffer without salt, incubated on ice for an additional 10 minutes, followed by centrifugation at maximal speed for 15 minutes. The supernatant was collected as soluble nucleolar fraction for IP analysis.

### Chromatin Fractionation

Soluble and chromatin fractions were extracted as previously described^30^. Cells were lysed in 400μL EBC buffer A consisting of 50mM Tris-HCl, pH 7.5, 100mM NaCl, 1mM EDTA, 2μg/mL aprotinin, 20mM NEM, 1mM PMSF, 1mM DTT, 0.05% NP40 for 8 minutes, followed by centrifugation at 800g for 5 minutes. The supernatants were collected as the soluble fractions. The pellets were then washed with EBC buffer A twice and a final wash with cold PBS. The pellets were then resuspended in 120 µL of EBC B buffer consisting of 50mM Tris-HCl, pH 7.5, 300mM NaCl, 0.5 mM EDTA, 2μg/mL aprotinin, 20mM NEM, 1mM PMSF, 5mM CaCl2, 50U micrococcal nuclease and incubated at room temperature for 10 minutes. The samples were then sonicated and centrifuged at 18,000 g at 4°C for 15 minutes. The supernatants were collected as the chromatin fractions.

### Gene knockdown by RNA interference

For siRNA-mediated knockdown, the 21-nucleotide siRNA duplexes with a 3’ dTdT overhang were synthesized by Dharmacon Inc (Lafayette, CO). The target sequences for GNL3 were 5’- GAACTAAAACAGCAGCAGA-3’ (si1) and 5’-AAGCTGTACTGCCAAGAAC-3’ (si2). The target sequences for SENP3 were 5’-ACTCCGTACCAAGGGTTAT-3’ (si1) and 5’- CTGGCCCTGTCTCAGCCAT-3’ (si2). The target sequence for USP36 was 5’- TGTCCTGAGTGGAGAGAAT-3’. The control scramble RNA was previously described^98^. These siRNA duplexes (100 nM) were introduced into cells using Lipofectamine 2000 (Invitrogen) following the manufacturer’s protocol. Cells were harvested at 48-72 hours post-transfection.

### Comet assay

Cells were trypsinized, pelleted at 200g for 5 minutes, and resuspended in PBS. The cell suspension was mixed with molten 1% low melting agarose (LMA, 1:10 ratio), spread on glass slides coated with a 1% normal melting point agarose (NMA) layer, solidified at 4°C for 10 minutes, and lysed with lysis buffer (2.5 M NaCl, 100 mM EDTA, 10 mM Tris base, 200 mM NaOH, 1% sodium lauryl sarcosinate, 1% Triton X-100) at 4°C overnight in the dark. Slides were equilibrated in neutral electrophoresis buffer (100 mM Tris, 300 mM sodium acetate, pH 9.0) at 4°C for 30 minutes and then electrophoresed at 1 V/cm (20 V, 400 mA) for 45 minutes. Slides were then sequentially dehydrated in 80% and 100% ethanol, 5 minutes each, respectively, and stained with SYBR Green (1:2000 in PBS, pH 7.4) for 15 minutes. After rinsing, slides were dried and imaged immediately. Tail moment was quantified from the images using ImageJ software.

### In vivo ubiquitination and SUMOylation assays

In vivo ubiquitination and SUMOylation assays under denaturing conditions were conducted using a Ni^2+^-NTA pull-down (PD) method as previously described^66^. For ubiquitination assay, cells were transfected with His-Ub and indicated plasmids and treated with 20 μM MG132 for 6 hours before harvesting. The cells were harvested 48 hours after transfection. A total of 20% of the cells were used for direct IB, while the remaining cells were subjected to Ni^2+^-NTA PD under denaturing conditions. The bead-bound proteins were analyzed by IB. For SUMOylation assay, cells were transfected with His-SUMO1 or His-SUMO2 and indicated plasmids, followed by Ni^2+^- NTA under denaturing conditions as above.

### Affinity purification and Mass spectrometry

293 cells stably expressing control or Flag-GNL3 were lysed in NP40 lysis buffer supplemented with protease inhibitors at 4 °C for one hour followed by centrifugation. Twenty milligrams of the cleared cell lysates from either control or Flag-GNL3 expressing cells were incubated with 0.1 ml of anti-Flag (M2) agarose beads at 4°C for 4 hours. The beads were washed four times in lysis buffer containing protease inhibitors. The bead-bound proteins were eluted in 0.2 ml of TBS (50 mM Tris-HCl, 150 mM NaCl, pH 7.4) containing 0.1 mg/ml of Flag peptides. Eluted proteins were dried in a SpeedVac and resuspended in SDS protein extraction buffer (5% SDS, 50 mM TEAB) and samples were solubilized in the detergent buffer by shaking at 90°C for 5 min. Samples were reduced by adding dithiothreitol and alkylated by the addition of iodoacetamide. Samples were then acidified by the addition of 12% aqueous phosphoric acid, and S-Trap protein binding buffer (90% aqueous methanol containing a final concentration of 100 mM TEAB, pH 8) was added. The acidified SDS lysate/MeOH S-Trap buffer mixture from each sample was transferred to the S-trap micro column (Protifi, New York). Samples were centrifuged until all SDS lysate/S-Trap buffer had passed through the S-Trap column. Each S-Trap micro column was washed 5 times by adding the S-Trap protein binding buffer (90% aqueous methanol with 100mM TEAB). Sequencing grade modified trypsin (Promega, Fitchburg, WI, Cat # V5111) was added to the S-trap columns and digestion was performed at 37°C overnight in a humidified chamber. Upon completion of digestion, peptides were eluted by sequential addition and centrifugation of 100 mM TEAB, 0.2% aqueous formic acid and 50%/50% acetonitrile/0.2% formic acid. The elution fractions were combined and then dried by vacuum centrifugation. Samples were reconstituted with HPLC grade water at 37°C in a shaker and peptide concentrations were determined using a Pierce Quantitative Colorimetric Peptide Assay (ThermoFisher Scientific, Cat # 23275). Recovered peptides were then dried by vacuum centrifugation, dissolved in 5% formic acid, loaded onto an Acclaim PepMap 0.1 x 20 mm NanoViper C18 peptide trap (Thermo Scientific) for 5 min at 10 µl/min in a 2% acetonitrile (ACN), 0.1% formic acid mobile phase. Peptides were then separated using a PepMap RSLC C18, 2- micron particle, 75-micron x 50-cm EasySpray column using a 7.5–30% ACN gradient over 90 min in mobile phase containing 0.1% formic acid and a 300 nl/min flow rate provided by a Dionex NCS-3500RS UltiMate RSLC nano UPLC (Thermo Scientific). Tandem mass spectrometry data was collected using an Orbitrap Q-Exactive mass spectrometer (Thermo Scientific) configured for top 10 data dependent analysis. MS1 scan resolution was set to 120,000 (at m/z 200) and MS1 automatic gain control (AGC) target was 3,000,000 with a maximum injection time of 50ms. Mass range was set at 375-1400. MS1 data were acquired in profile mode using positive polarity, MIPS filter on with relaxed conditions, and charge states from +2 to +7 accepted. The Orbitrap AGC target value for centroid-mode fragment spectra was set at 100,000 (max IT 100ms) and the intensity threshold was 50,000. Quadrupole isolation width was set at 1.2 m/z. Normalized collision energy was set at 30%. The program Comet (v. 2016.01, rev. 3) was used to search MS2 Spectra against an April 2024 version of a UniProt FASTA protein database containing 20,598 canonical Homo sapiens sequences and 175 common contaminant sequences. To estimate error rates, sequence-reversed forms of all proteins were concatenated to the FASTA file. The database processing was performed with Python scripts available at https://github.com/pwilmart/fasta_utilities.git and Comet results processing used the PAW pipeline^99^ from https://github.com/pwilmart/PAW_pipeline.git. Comet searches for all samples were performed with trypsin enzyme specificity. Monoisotopic parent ion mass tolerance was 1.25 Da. Monoisotopic fragment ion mass tolerance was 0.02 Da. A static modification of +57.021464 Da was added to all cysteine residues and a variable modification of +15.9949 Da to methionine residues. Comet scores were combined into linear discriminant function scores, and discriminant score histograms were created separately for each peptide charge state (2+, 3+, and 4+). Separate histograms were created for matches to forward sequences and for matches to reversed sequences for all peptides of 7 amino acids or longer. The score histograms for reversed matches were used to estimate peptide false discovery rates (FDR) and set score thresholds for each peptide class. After removal of contaminates, this resulted in the identification of 1,526 proteins with at least two unique peptides per protein and an estimated protein false discovery rate <0.4%. Estimation of protein abundance differences was performed using the numbers of assigned MS/MS spectra (spectral counts) to each peptide from protein across the two samples. Statistical analysis was performed using Excel (Microsoft, WA) to perform a Student’s T-test to compare the spectral counts for each protein between groups.

### Cell viability and Colony formation assays

Cell viability was measured by 3-(4,5-dimethylthiazol-2-yl)-2,5-diphenyltetrazolium bromide (MTT) assays. Briefly, cells were incubated with 0.5 mg/ml MTT in medium for 3 hours. After incubation, MTT medium was removed and dimethylsulfoxide (DMSO, 100 μl per well) was added to fully dissolve the purple formazan. The absorbance was measured at OD560nm and OD690nm. The reduced absorbance (Abs560nm-Abs690nm) represents the relative number of viable cells per well. Colony formation was used to measure cell proliferation. Cells were seeded into 60 mm plates at a density of 2,500 cells per plate and cultured in DMEM medium containing 10% FBS for up to 2 weeks. The colonies were visualized by staining with 0.5% crystal violet in 50% ethanol at room temperature for 2 h. Captured digital images of colonies were further processed and quantified using the ImageJ software.

### Proximity Ligation Assay

Proximity ligation assays were then performed using the Duolink® PLA Fluorescence Kit (Sigma-Aldrich) according to the manufacturer’s instructions, with minor modifications. Briefly, cells cultured in cover slides were fixed with 4% paraformaldehyde for 15 minutes, permeabilized with 0.25% Triton X-100 for 15 minutes and blocked with Duolink® blocking solution at 37 °C for 1 hour. The slides were then incubated with primary antibodies at 4 °C overnight. After washing with Duolink® In Situ Wash Buffer A, PLUS and MINUS PLA probes were diluted 1:5 in Duolink® antibody diluent buffer and applied to the slides, followed by incubation at 37 °C for 1 hour. The slides were then washed with Wash Buffer A, and ligation was performed by adding ligase to 1× ligation buffer at a 1:40 dilution, followed by incubation in a pre-heated humidity chamber at 37 °C for 30 minutes. After additional washing with Wash Buffer A, amplification was carried out by adding polymerase to 1× amplification buffer at a 1:80 dilution, followed by incubation and subsequent washing with Wash Buffer B. Finally, the slides were mounted with Duolink® In Situ Mounting Medium containing DAPI and covered with coverslips. Samples were allowed to equilibrate for 15 minutes before imaging with a confocal microscope.

### Immunofluorescence staining

Cells cultured in cover slides were fixed with 4% paraformaldehyde, permeabilized with 0.25% Triton X-100, and then blocked with 8% BSA. The cells were stained with primary antibodies followed by staining with Alexa Fluor 488 (green) goat anti-mouse, Alexa Fluor 555 (red) goat anti-mouse, Alexa Fluor 488 (green) goat anti-rabbit, or Alexa Fluor 555 (red) goat anti-rabbit antibodies (Invitrogen) as well as DAPI for DNA staining (see listed antibodies in Supplementary Table 2). Stained cells were imaged by using a Zeiss ApTome fluorescence microscope (Zeiss) or a Zeiss LSM980 confocal microscope (Zeiss).

### Image analysis

Confocal imaging was performed on a Zeiss LSM980 microscope equipped with a Plan-Apochromat 63 x/1.40 NA oil immersion objective lens (Zeiss). The following excitation and detection windows were used: DAPI, 405 nm excitation, 420–480 nm emission; Alexa Fluor 488, 488 nm excitation, 500–550 nm emission; Alexa Fluor 555, 561 nm excitation, 565–620 nm emission. PLA signals were quantified as the number of discrete nuclear dots per cell using ImageJ. DNA damage-induced foci were also quantified using ImageJ. For all PLA and DNA damage foci experiments, at least 100 cells were counted from at least 3 independent experiments.

### DNA End resection assay

Cells were seeded in 24-well plates at a density of 3-4x10^4^ cells/well and Cells were cultured in coverslips in the presence of 10 μM BrdU for 20 hours. Cells were washed twice with PBS and incubated in pre-extraction buffer A (10 mM pH 7 PIPES, 100 mM NaCl, 3 mM MgCl_2_, 1 mM EGTA, 0.5% Triton X-100 and 300 mM Sucrose) at 4°C for 10 minutes. Then cells were incubated in cytoskeleton stripping buffer B (10 mM Tris pH 7.5, 10 mM NaCl, 3 mM MgCl_2_, 1% Tween 20, 0.50% sodium deoxycholate) at 4°C for 10 minutes and washed once with PBS. Cells were fixed with 4% paraformaldehyde for 15 minutes and followed by 100% cold methanol at -20°C for 5 minutes. Cells were permeabilized with 0.5% Triton X-100 at room temperature for 15 minutes, blocked in 3% BSA, and incubated with anti-BrdU and anti-GNL3 antibodies, followed by staining with Alexa Fluor 488 (green) goat anti-mouse, Alexa Fluor 555 (red) goat anti-rabbit antibodies as well as DAPI for DNA staining. BrdU-positive cells were defined as nuclei containing ≥ 20 strong BrdU foci based on previously published criteria ^100,101^.

### PLA-Based Detection of Protein Association with ssDNA

Cells were labeled with 10 μM BrdU as described in the End Resection Assay. After fixation and permeabilization, PLA was performed using the Duolink® In Situ Red Starter Kit (Sigma-Aldrich) with anti-BrdU and anti-GNL3 antibodies following the manufacturer’s instructions. Slides were mounted using Duolink® In Situ Mounting Medium with DAPI and imaged using Zeiss LSM980 confocal microscope (Zeiss).

### HR and NHEJ reporter assays

U2OS Cells were transfected with scr or SENP3 siRNA together with pCBA-I-SceI, pCherry, and either DR-GFP or SSA-GFP plasmids. The cells were harvested and fixed at 48 hours after transection and analyzed by flowcytometry for the percentage of mCherry/GFP positive cells.

The expression of mCherry was used for the normalization of transfection efficiency.

### TCGA data analysis

RNA-seq and clinical data from The Cancer Genome Atlas (TCGA) were retrieved using the R package TCGAbiolinks (v2.30.4). Gene-level expression was quantified as transcripts per million (TPM) and transformed as log1p(TPM) prior to analysis. Triple-negative breast cancer (TNBC) status was defined from TCGA receptor annotations for ER, PR, and HER2 negative; cases missing any receptor annotation were excluded. Group differences were assessed with two-sided Wilcoxon rank-sum tests.

### Statistical analysis

Measurements and cell counts were performed using ImageJ. Results are expressed as the mean ± s.d. Statistical analysis was performed using GraphPad Prism. Unpaired two-tailed Student’s test was employed to determine the differences between two groups. Two-way ANOVA was employed to determine statistical differences among multiple groups, followed by Bonferroni’s multiple comparisons test. The level of statistical significance stated in the text is based on p-values. p < 0.05 was considered statistically significant.

## ACKNOWLEDGMENTS

We thank Dr. Masayuki Komada (Tokyo Institute of Technology, Japan) for providing anti-USP36 antisera, Dr. Zhenkun Lou (Mayo Clinic) for providing plasmids, Phillip Wilmarth for helping with PRIDE deposition, and Dr. Gabriel Cohn for technical assistance. This work was supported by grants from NIH grants R01 CA262104 and R35 GM153360 to M-S. D. Mass spectrometric analysis was performed by the OHSU Proteomics Shared Resource (RRID: SCR_009991) with partial support from NIH core grants P30EY010572, P30CA069533, and OHSU Emerging Technology Fund.

## Author contributions

MSD and YY conceived and designed experiments; YY, YL, RSD, XXS and MSD performed the experiments and/or analyzed the data. CC and ZX performed TCGA data analysis. KZ and AR performed proteomic analysis. RCS provided discussion for this study. MSD and XXS supervised the research. YY and MSD wrote the manuscript.

## Conflict of Interest

The authors declare no competing interests.

## Data Availability

The mass spectrometry proteomics data generated in this study were deposited to the ProteomeXchange Consortium via the PRIDE partner repository with the dataset identifier PXD068465 (https://www.ebi.ac.uk/pride/). (Reviewer account details: Username: Password: wzHH2Um8sG1K). All data generated or analyzed in this study are included in this article and its Supplementary Information files.

## Notes

### Competing Interest Statement

The authors have declared no competing interest.

## References

1 Jackson, S. P. & Bartek, J. The DNA-damage response in human biology and disease. Nature 461, 1071–1078 (2009).

2 Ciccia, A. & Elledge, S. J. The DNA damage response: making it safe to play with knives. Mol Cell 40, 179–204 (2010).

3 Chang, H. H. Y., Pannunzio, N. R., Adachi, N. & Lieber, M. R. Non-homologous DNA end joining and alternative pathways to double-strand break repair. Nature reviews. Molecular cell biology 18, 495–506 (2017).

4 Hustedt, N. & Durocher, D. The control of DNA repair by the cell cycle. Nat Cell Biol 19, 1–9 (2016).

5 Scully, R., Panday, A., Elango, R. & Willis, N. A. DNA double-strand break repair-pathway choice in somatic mammalian cells. Nat Rev Mol Cell Biol 20, 698–714 (2019).

6 Sfeir, A., Tijsterman, M. & McVey, M. Microhomology-Mediated End-Joining Chronicles: Tracing the Evolutionary Footprints of Genome Protection. Annu Rev Cell Dev Biol 40, 195–218 (2024).

7 Zhao, F., Kim, W., Kloeber, J. A. & Lou, Z. DNA end resection and its role in DNA replication and DSB repair choice in mammalian cells. Exp Mol Med 52, 1705–1714 (2020).

8 Ceccaldi, R. & Cejka, P. Mechanisms and regulation of DNA end resection in the maintenance of genome stability. Nat Rev Mol Cell Biol 26, 586–599 (2025).

9 Stracker, T. H. & Petrini, J. H. The MRE11 complex: starting from the ends. Nat Rev Mol Cell Biol 12, 90–103 (2011).

10 Groelly, F. J., Fawkes, M., Dagg, R. A., Blackford, A. N. & Tarsounas, M. Targeting DNA damage response pathways in cancer. Nat Rev Cancer 23, 78–94 (2023).

11 Foo, T. K. & Xia, B. BRCA1-Dependent and Independent Recruitment of PALB2-BRCA2- RAD51 in the DNA Damage Response and Cancer. Cancer Res 82, 3191–3197 (2022).

12 Holloman, W. K. Unraveling the mechanism of BRCA2 in homologous recombination. Nat Struct Mol Biol 18, 748–754 (2011).

13 Guo, M. & Wang, S. M. The BRCAness Landscape of Cancer. Cells 11 (2022).

14 Curtin, N. J. & Szabo, C. Poly(ADP-ribose) polymerase inhibition: past, present and future. Nat Rev Drug Discov 19, 711–736 (2020).

15. Jackson, L. M. & Moldovan, G. L. Mechanisms of PARP1 inhibitor resistance and their implications for cancer treatment. NAR Cancer 4, zcac042 (2022).

16 Fu, X. et al. Mechanism of PARP inhibitor resistance and potential overcoming strategies. Genes Dis 11, 306–320 (2024).

17 Geiss-Friedlander, R. & Melchior, F. Concepts in sumoylation: a decade on. Nat Rev Mol Cell Biol 8, 947–956 (2007).

18 Vertegaal, A. C. O. Signalling mechanisms and cellular functions of SUMO. Nat Rev Mol Cell Biol 23, 715–731 (2022).

19 Gareau, J. R. & Lima, C. D. The SUMO pathway: emerging mechanisms that shape specificity, conjugation and recognition. Nat Rev Mol Cell Biol 11, 861–871 (2010).

20 Dou, H., Huang, C., Van Nguyen, T., Lu, L. S. & Yeh, E. T. SUMOylation and de- SUMOylation in response to DNA damage. FEBS Lett 585, 2891–2896 (2011).

21 Gasser, S. M. & Stutz, F. SUMO in the regulation of DNA repair and transcription at nuclear pores. FEBS Lett 597, 2833–2850 (2023).

22 Jackson, S. P. & Durocher, D. Regulation of DNA damage responses by ubiquitin and SUMO. Mol Cell 49, 795–807 (2013).

23 Schwertman, P., Bekker-Jensen, S. & Mailand, N. Regulation of DNA double-strand break repair by ubiquitin and ubiquitin-like modifiers. Nat Rev Mol Cell Biol 17, 379–394 (2016).

24 Galanty, Y. et al. Mammalian SUMO E3-ligases PIAS1 and PIAS4 promote responses to DNA double-strand breaks. Nature 462, 935–939 (2009).

25 Morris, J. R. et al. The SUMO modification pathway is involved in the BRCA1 response to genotoxic stress. Nature 462, 886–890 (2009).

26 Luo, K., Zhang, H., Wang, L., Yuan, J. & Lou, Z. Sumoylation of MDC1 is important for proper DNA damage response. EMBO J 31, 3008–3019 (2012).

27 Galanty, Y., Belotserkovskaya, R., Coates, J. & Jackson, S. P. RNF4, a SUMO-targeted ubiquitin E3 ligase, promotes DNA double-strand break repair. Genes Dev 26, 1179–1195 (2012).

28 Vyas, R. et al. RNF4 is required for DNA double-strand break repair in vivo. Cell Death Differ 20, 490–502 (2013).

29 Yin, Y. et al. SUMO-targeted ubiquitin E3 ligase RNF4 is required for the response of human cells to DNA damage. Genes Dev 26, 1196–1208 (2012).

30 Zhang, T. et al. Crosstalk between SUMOylation and ubiquitylation controls DNA end resection by maintaining MRE11 homeostasis on chromatin. Nat Commun 13, 5133 (2022).

31 Ouyang, K. J. et al. SUMO modification regulates BLM and RAD51 interaction at damaged replication forks. PLoS Biol 7, e1000252 (2009).

32 Eladad, S. et al. Intra-nuclear trafficking of the BLM helicase to DNA damage-induced foci is regulated by SUMO modification. Hum Mol Genet 14, 1351–1365 (2005).

33 Ouyang, K. J., Yagle, M. K., Matunis, M. J. & Ellis, N. A. BLM SUMOylation regulates ssDNA accumulation at stalled replication forks. Front Genet 4, 167 (2013).

34 Claessens, L. A., Verlaan-de Vries, M., de Graaf, I. J. & Vertegaal, A. C. O. SENP6 regulates localization and nuclear condensation of DNA damage response proteins by group deSUMOylation. Nat Commun 14, 5893 (2023).

35 Kumar, R., Gonzalez-Prieto, R., Xiao, Z., Verlaan-de Vries, M. & Vertegaal, A. C. O. The STUbL RNF4 regulates protein group SUMOylation by targeting the SUMO conjugation machinery. Nat Commun 8, 1809 (2017).

36 Dou, H., Huang, C., Singh, M., Carpenter, P. B. & Yeh, E. T. Regulation of DNA repair through deSUMOylation and SUMOylation of replication protein A complex. Mol Cell 39, 333–345 (2010).

37 Antoniuk-Majchrzak, J. et al. Stability of Rad51 recombinase and persistence of Rad51 DNA repair foci depends on post-translational modifiers, ubiquitin and SUMO. Biochim Biophys Acta Mol Cell Res 1870, 119526 (2023).

38 Hariharasudhan, G. et al. TOPORS-mediated RAD51 SUMOylation facilitates homologous recombination repair. Nucleic Acids Res 50, 1501–1516 (2022).

39 Shima, H. et al. Activation of the SUMO modification system is required for the accumulation of RAD51 at sites of DNA damage. J Cell Sci 126, 5284–5292 (2013).

40 Altmannova, V. et al. Rad52 SUMOylation affects the efficiency of the DNA repair. Nucleic Acids Res 38, 4708–4721 (2010).

41 Sacher, M., Pfander, B., Hoege, C. & Jentsch, S. Control of Rad52 recombination activity by double-strand break-induced SUMO modification. Nat Cell Biol 8, 1284–1290 (2006).

42 Torres-Rosell, J. et al. The Smc5-Smc6 complex and SUMO modification of Rad52 regulates recombinational repair at the ribosomal gene locus. Nat Cell Biol 9, 923–931 (2007).

43 Whalen, J. M., Dhingra, N., Wei, L., Zhao, X. & Freudenreich, C. H. Relocation of Collapsed Forks to the Nuclear Pore Complex Depends on Sumoylation of DNA Repair Proteins and Permits Rad51 Association. Cell Rep 31, 107635 (2020).

44 Li, Y. J., Stark, J. M., Chen, D. J., Ann, D. K. & Chen, Y. Role of SUMO:SIM-mediated protein-protein interaction in non-homologous end joining. Oncogene 29, 3509–3518 (2010).

45 Guzzo, C. M. et al. RNF4-dependent hybrid SUMO-ubiquitin chains are signals for RAP80 and thereby mediate the recruitment of BRCA1 to sites of DNA damage. Sci Signal 5, ra88 (2012).

46 Chang, Y. C., Oram, M. K. & Bielinsky, A. K. SUMO-Targeted Ubiquitin Ligases and Their Functions in Maintaining Genome Stability. Int J Mol Sci 22 (2021).

47 Groocock, L. M. et al. RNF4 interacts with both SUMO and nucleosomes to promote the DNA damage response. EMBO Rep 15, 601–608 (2014).

48 Heideker, J., Perry, J. J. & Boddy, M. N. Genome stability roles of SUMO-targeted ubiquitin ligases. DNA Repair (Amst) 8, 517–524 (2009).

49 Pfeiffer, A. et al. Ataxin-3 consolidates the MDC1-dependent DNA double-strand break response by counteracting the SUMO-targeted ubiquitin ligase RNF4. EMBO J 36, 1066–1083 (2017).

50 Prudden, J. et al. SUMO-targeted ubiquitin ligases in genome stability. EMBO J 26, 4089–4101 (2007).

51 Tsai, R. Y. & McKay, R. D. A nucleolar mechanism controlling cell proliferation in stem cells and cancer cells. Genes Dev 16, 2991–3003 (2002).

52 Romanova, L. et al. Critical role of nucleostemin in pre-rRNA processing. J Biol Chem 284, 4968–4977 (2009).

53 Vanden Broeck, A. & Klinge, S. Principles of human pre-60S biogenesis. Science 381, eadh3892 (2023).

54 Ma, H. & Pederson, T. Depletion of the nucleolar protein nucleostemin causes G1 cell cycle arrest via the p53 pathway. Mol Biol Cell 18, 2630–2635 (2007).

55 Dai, M. S., Sun, X. X. & Lu, H. Aberrant expression of nucleostemin activates p53 and induces cell cycle arrest via inhibition of MDM2. Mol Cell Biol 28, 4365–4376 (2008).

56 Meng, L., Lin, T. & Tsai, R. Y. Nucleoplasmic mobilization of nucleostemin stabilizes MDM2 and promotes G2-M progression and cell survival. J Cell Sci 121, 4037–4046 (2008).

57 Huang, G., Meng, L. & Tsai, R. Y. p53 Configures the G2/M Arrest Response of Nucleostemin-Deficient Cells. Cell Death Discov 1, 15060- (2015).

58 Kobayashi, T. et al. Nucleostemin expression in invasive breast cancer. BMC Cancer 14, 215 (2014).

59 Yoshida, R. et al. Overexpression of nucleostemin contributes to an advanced malignant phenotype and a poor prognosis in oral squamous cell carcinoma. Br J Cancer 111, 2308–2315 (2014).

60 Hua, L. et al. Upregulated expression of Nucleostemin/GNL3 is associated with poor prognosis and Sorafenib Resistance in Hepatocellular Carcinoma. Pathol Res Pract 213, 688–697 (2017).

61 Sami, M. M., Hachim, M. Y., Hachim, I. Y., Elbarkouky, A. H. & Lopez-Ozuna, V. M. Nucleostemin expression in breast cancer is a marker of more aggressive phenotype and unfavorable patients’ outcome: A STROBE-compliant article. Medicine (Baltimore) 98, e14744 (2019).

62 Wang, J. et al. Nucleostemin Modulates Outcomes of Hepatocellular Carcinoma via a Tumor Adaptive Mechanism to Genomic Stress. Mol Cancer Res 18, 723–734 (2020).

63 Meng, L. et al. Nucleostemin deletion reveals an essential mechanism that maintains the genomic stability of stem and progenitor cells. Proc Natl Acad Sci U S A 110, 11415–11420 (2013).

64 Lin, T. et al. Nucleostemin reveals a dichotomous nature of genome maintenance in mammary tumor progression. Oncogene 38, 3919–3931 (2019).

65 Lebdy, R. et al. The nucleolar protein GNL3 prevents resection of stalled replication forks. EMBO Rep 24, e57585 (2023).

66 Ryu, H. et al. The deubiquitinase USP36 promotes snoRNP group SUMOylation and is essential for ribosome biogenesis. EMBO Rep 22, e50684 (2021).

67 Chen, Y. et al. The ubiquitin-specific protease USP36 SUMOylates EXOSC10 and promotes the nucleolar RNA exosome function in rRNA processing. Nucleic Acids Res (2023).

68 Li, Y. et al. The Ubiquitin-specific Protease USP36 Associates with the Microprocessor Complex and Regulates miRNA Biogenesis by SUMOylating DGCR8. Cancer Res Commun 3, 459–470 (2023).

69 Li, Y., Yang, Y., Sears, R. C., Dai, M. S. & Sun, X. X. USP36 SUMOylates Las1L and Promotes Its Function in Pre-Ribosomal RNA ITS2 Processing. Cancer Res Commun 4, 2835–2845 (2024).

70 Yang, Y., Li, Y., Sears, R. C., Sun, X. X. & Dai, M. S. SUMOylation regulation of ribosome biogenesis: Emerging roles for USP36. Front RNA Res 2 (2024).

71 Bhachoo, J. S. & Garvin, A. J. SUMO and the DNA damage response. Biochem Soc Trans 52, 773–792 (2024).

72 Hande, K. R. Etoposide: four decades of development of a topoisomerase II inhibitor. Eur J Cancer 34, 1514–1521 (1998).

73 Baldwin, E. L. & Osheroff, N. Etoposide, topoisomerase II and cancer. Curr Med Chem Anticancer Agents 5, 363–372 (2005).

74 Zhao, Q. et al. GPS-SUMO: a tool for the prediction of sumoylation sites and SUMO-interaction motifs. Nucleic Acids Res 42, W325–330 (2014).

75 Mukherjee, B., Tomimatsu, N. & Burma, S. Immunofluorescence-based methods to monitor DNA end resection. Methods Mol Biol 1292, 67–75 (2015).

76 Lascorz, J., Codina-Fabra, J., Reverter, D. & Torres-Rosell, J. SUMO-SIM interactions: From structure to biological functions. Semin Cell Dev Biol 132, 193–202 (2022).

77 Zhu, J. et al. Small ubiquitin-related modifier (SUMO) binding determines substrate recognition and paralog-selective SUMO modification. J Biol Chem 283, 29405–29415 (2008).

78 Meng, L., Yasumoto, H. & Tsai, R. Y. Multiple controls regulate nucleostemin partitioning between nucleolus and nucleoplasm. J Cell Sci 119, 5124–5136 (2006).

79 Haindl, M., Harasim, T., Eick, D. & Muller, S. The nucleolar SUMO-specific protease SENP3 reverses SUMO modification of nucleophosmin and is required for rRNA processing. EMBO Rep 9, 273–279 (2008).

80 Yun, C. et al. Nucleolar protein B23/nucleophosmin regulates the vertebrate SUMO pathway through SENP3 and SENP5 proteases. J Cell Biol 183, 589–595 (2008).

81. Han, Y., et al. SENP3-mediated TIP60 deSUMOylation is required for DNA-PKcs activity and DNA damage repair. MedComm (2020) 3, e123 (2022).

82 Hu, G. et al. Mitotic SENP3 activation couples with cGAS signaling in tumor cells to stimulate anti-tumor immunity. Cell Death Dis 13, 640 (2022).

83 Xu, R. et al. hCINAP regulates the DNA-damage response and mediates the resistance of acute myelocytic leukemia cells to therapy. Nat Commun 10, 3812 (2019).

84 Brosh, R. M., Jr., et al. Replication protein A physically interacts with the Bloom’s syndrome protein and stimulates its helicase activity. J Biol Chem 275, 23500–23508 (2000).

85 Daley, J. M. et al. Enhancement of BLM-DNA2-Mediated Long-Range DNA End Resection by CtIP. Cell Rep 21, 324–332 (2017).

86 Clarke, T. L. et al. ZNF280A links DNA double-strand break repair to human 22q11.2 distal deletion syndrome. Nat Cell Biol 27, 1006–1020 (2025).

87 Bythell-Douglas, R. & Deans, A. J. A Structural Guide to the Bloom Syndrome Complex. Structure 29, 99–113 (2021).

88 Ismail, I. H. et al. CBX4-mediated SUMO modification regulates BMI1 recruitment at sites of DNA damage. Nucleic Acids Res 40, 5497–5510 (2012).

89 Bermudez-Lopez, M. et al. Sgs1’s roles in DNA end resection, HJ dissolution, and crossover suppression require a two-step SUMO regulation dependent on Smc5/6. Genes Dev 30, 1339–1356 (2016).

90 Bonner, J. N. et al. Smc5/6 Mediated Sumoylation of the Sgs1-Top3-Rmi1 Complex Promotes Removal of Recombination Intermediates. Cell Rep 16, 368–378 (2016).

91 Li, S. et al. Multifaceted regulation of the sumoylation of the Sgs1 DNA helicase. J Biol Chem 298, 102092 (2022).

92 Bugreev, D. V., Mazina, O. M. & Mazin, A. V. Bloom syndrome helicase stimulates RAD51 DNA strand exchange activity through a novel mechanism. J Biol Chem 284, 26349–26359 (2009).

93 Beekman, C. et al. Evolutionarily conserved role of nucleostemin: controlling proliferation of stem/progenitor cells during early vertebrate development. Mol Cell Biol 26, 9291–9301 (2006). h

94 Zhu, Q., Yasumoto, H. & Tsai, R. Y. Nucleostemin delays cellular senescence and negatively regulates TRF1 protein stability. Mol Cell Biol 26, 9279–9290 (2006).

95 Rubbi, C. P. & Milner, J. Disruption of the nucleolus mediates stabilization of p53 in response to DNA damage and other stresses. Embo J 22, 6068–6077 (2003).

96 Ogawa, L. M. & Baserga, S. J. Crosstalk between the nucleolus and the DNA damage response. Mol Biosyst 13, 443–455 (2017).

97 Meng, L., Hsu, J. K., Zhu, Q., Lin, T. & Tsai, R. Y. Nucleostemin inhibits TRF1 dimerization and shortens its dynamic association with the telomere. J Cell Sci 124, 3706–3714 (2011).

98 Dai, M. S. et al. Ribosomal protein L23 activates p53 by inhibiting MDM2 function in response to ribosomal perturbation but not to translation inhibition. Mol Cell Biol 24, 7654–7668 (2004).

99 Wilmarth, P. A., Riviere, M. A. & David, L. L. Techniques for accurate protein identification in shotgun proteomic studies of human, mouse, bovine, and chicken lenses. J Ocul Biol Dis Infor 2, 223–234 (2009).

100 Karlsson, K. H. & Stenerlow, B. Extensive ssDNA end formation at DNA double-strand breaks in non-homologous end-joining deficient cells during the S phase. BMC Mol Biol 8, 97 (2007).

101 Kurashima, K. et al. Poleta, a Y-family translesion synthesis polymerase, promotes cellular tolerance of Myc-induced replication stress. J Cell Sci 131 (2018).

