## Supplementary Figs 1-8 & Table 1-2 for "GNL3 SUMOylation is essential for DNA double-strand break repair by homologous recombination"

This PDF file includes:

Supplementary Figures 1 to 8

Supplementary Table 1-2

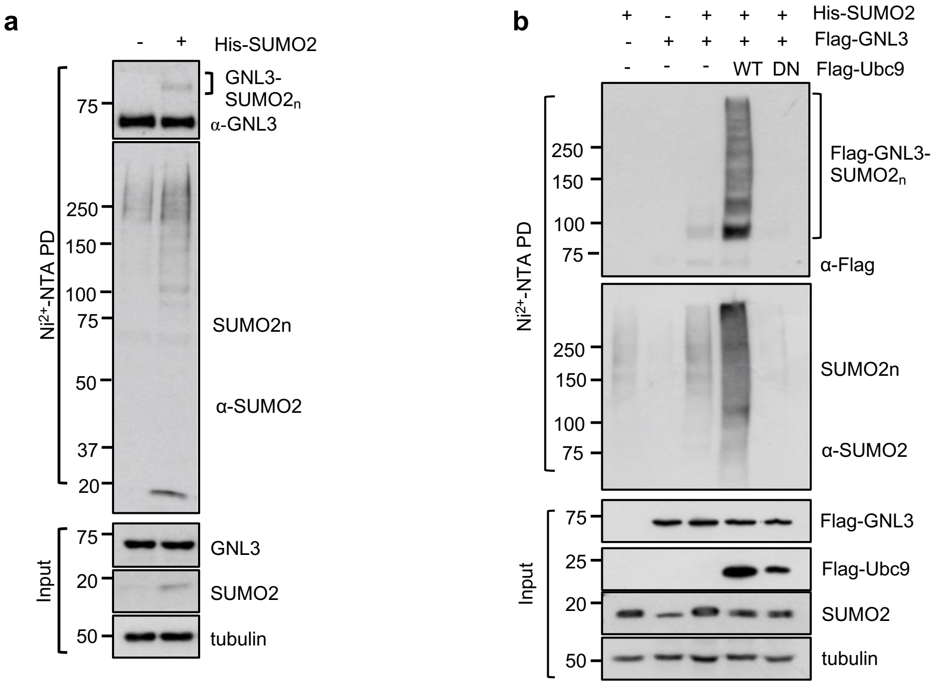


**Figure S1. DNA damage induces GNL3 SUMOylation. (a)** SUMOylation of endogenous GNL3 by SUMO2. HeLa cells transfected with control or His-SUMO2 were subjected to $Ni^{2+}$-NTA PD under denaturing conditions, followed by IB. **(b)** Overexpression of WT Ubc9, but not its dominant-negative (DN) mutant, promoted GNL3 SUMOylation. H1299 cells transfected with the indicated plasmids were subjected to $Ni^{2+}$-NTA PD under denaturing conditions, followed by IB. The SUMO2 modified GNL3 is indicated. The protein expression is shown at the bottom panels.

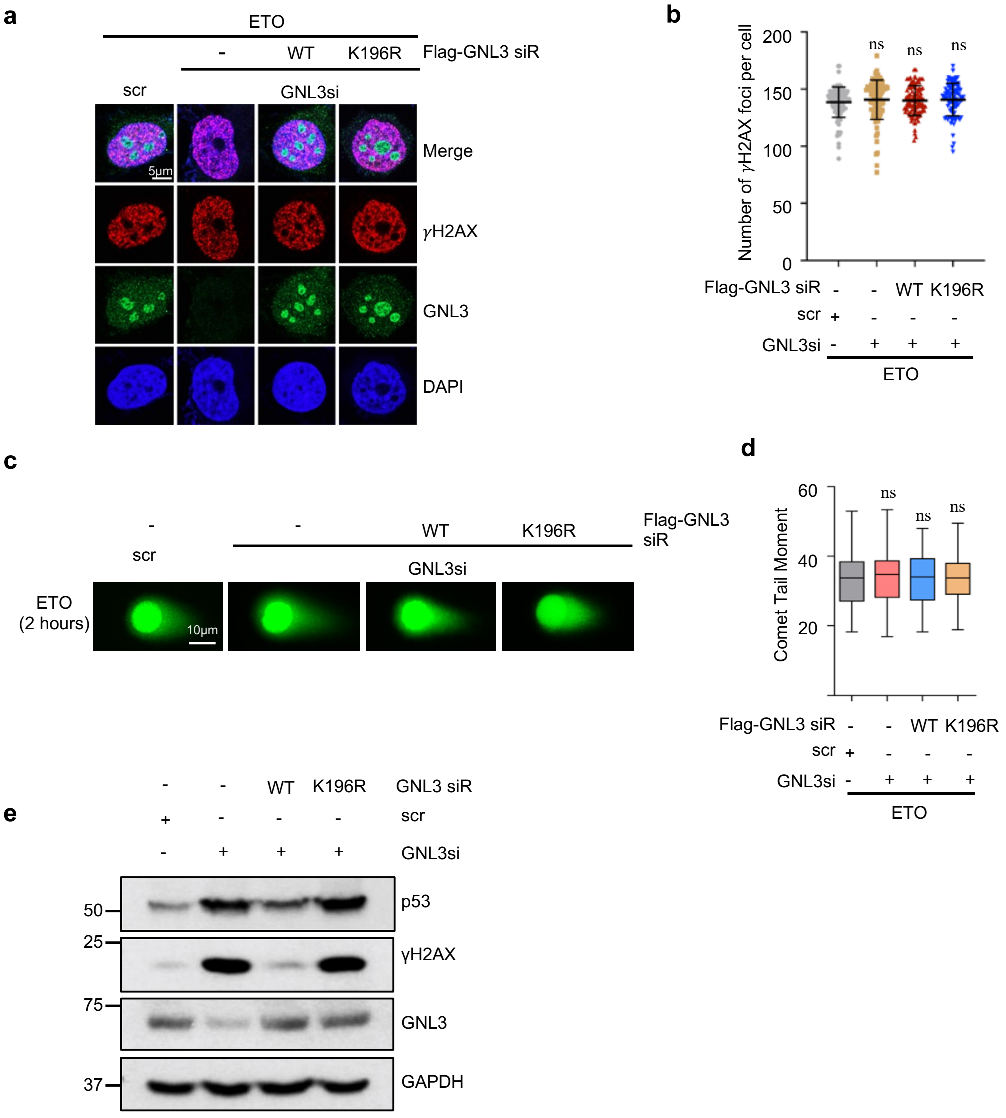


**Figure S2. GNL3 SUMOylation at K196 is essential for DDR. (a)(b)** Knockdown of GNL3 does not abolish DNA damage induced γH2AX foci. U2OS cells transfected with scrambled (scr) or GNL3 siRNA together with Flag-tagged siRNA-resistant (siR) WT GNL3 or the K196R mutant were treated with 20 μM ETO for 2 hours and assayed by IF staining with anti-GNL3 and anti-γH2AX antibodies. Shown are representative confocal images (a) and the quantification (b). **(c)(d)** DSB repair in GNL3 knockdown and rescue experiments upon ETO treatment. U2OS cells transfected with scr or GNL3 siRNA together with Flag-tagged siRNA-resistant WT GNL3 or the K196R mutant were treated with 20 μM ETO for 2 hours and subjected to Comet assays. Shown are representative images (c) and the quantification (d). **(e)** Ectopic expression of WT GNL3, but not the K196R mutant, abolishes p53 induction upon knockdown of endogenous GNL3. U2OS cells transfected with scrambled (scr) or GNL3 siRNA together with Flag-tagged siRNA-resistant (siR) WT GNL3 or the K196R mutant were assayed IB. **P<0.01, compared to controls. ns, not significant.

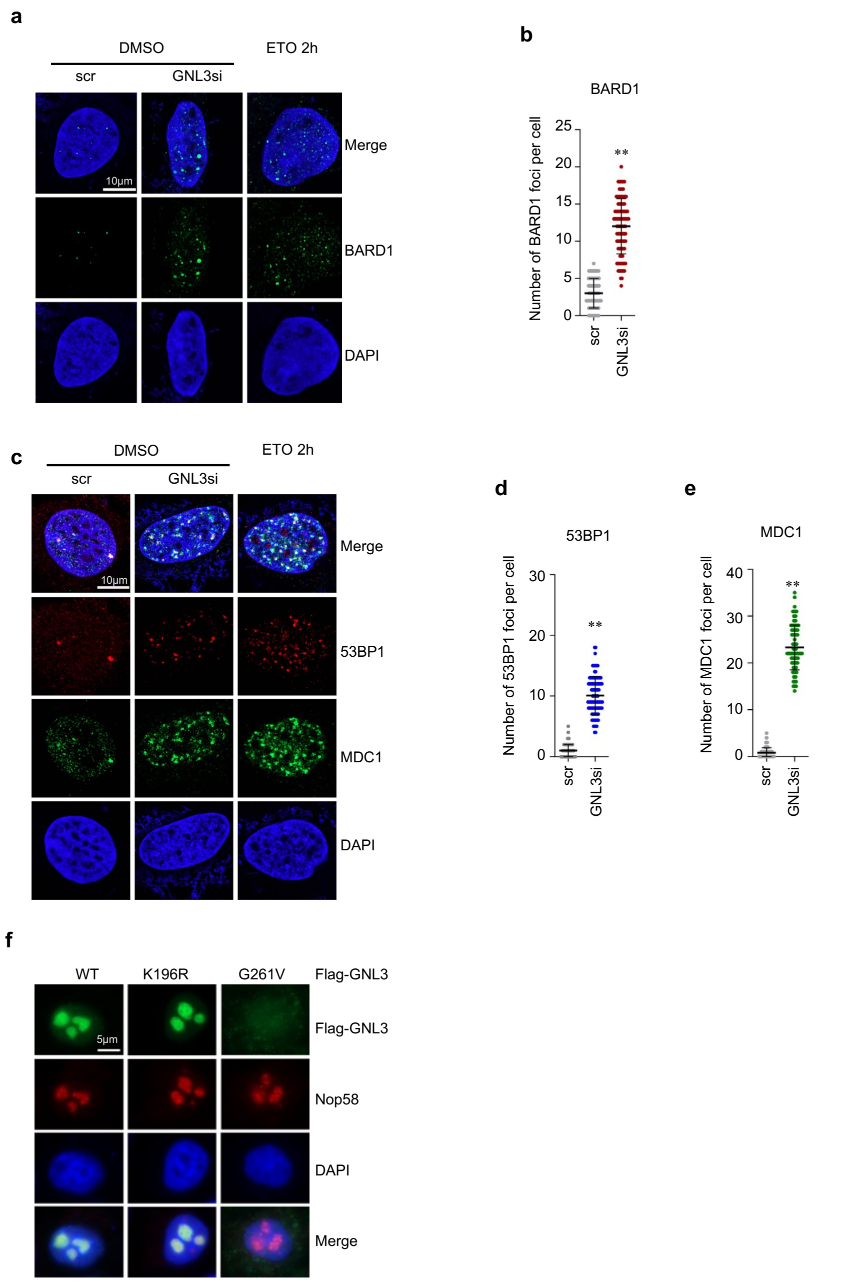


**Figure S3. GNL3 knockdown does not affect the foci formation of BARD1, 53BP1 and MDC1.** U2OS cells transfected with scr or GNL3 siRNA or treated with 20 μM ETO for 2 hours were assayed by IF using anti-BARD1 **(a)(b)** or anti-53BP1 and anti-MDC1 **(c)-(e).** Shown are representative confocal images (a)(c) and the quantification (b)(d)(e). **P<0.01, compared to control group. **(f)**The SUMO-defective K196R, but not the GTP-binding defective G261V mutant, is localized in the nucleolus**.** U2OS cells transfected with the indicated plasmids were assayed by IF staining with anti-Flag and anti-Nop58 (a nucleolar marker) antibodies.

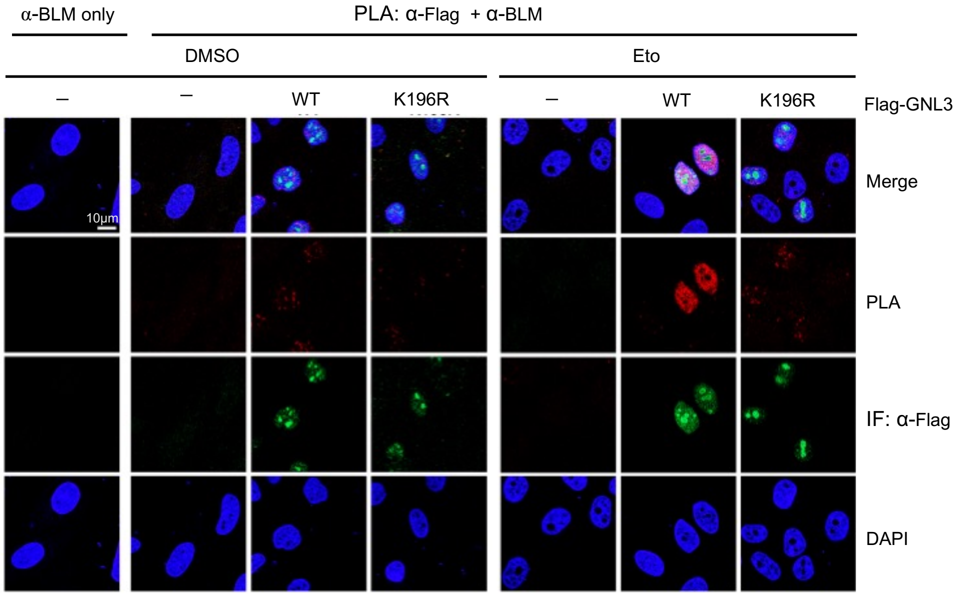


**Figure S4. GNL3 regulates DNA end resection by interacting with BLM-DNA2.** GNL3 SUMOylation at K196 is required for interaction with BLM upon DNA damage. U2OS cells transfected with WT GNL3 or the K196R mutant were treated with control or 20 μM ETO for 2 hours and assayed by PLA using anti-Flag and anti-BLM antibodies. Anti-BLM only staining is used as a control.

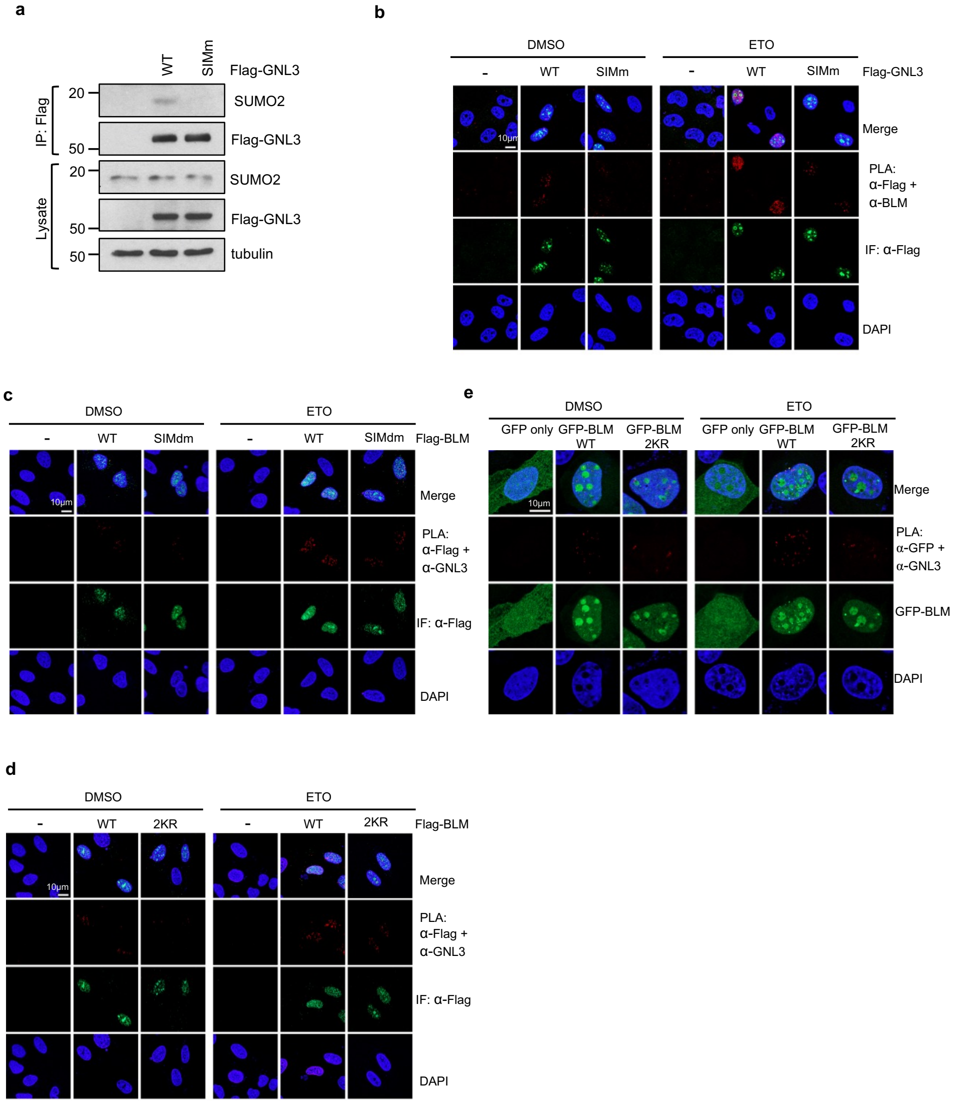


**Figure S5. SUMO-SIM interaction directs the GNL3 interaction with BLM. (a)** Mutating the SIM abolishes GNL3 interaction with SUMO2. H1299 cells transfected with WT GNL3 or the SIMm mutant were assayed by co-IP using anti-Flag antibody, followed by IB. **(b)** Mutating the SIM attenuates GNL3 interaction with BLM in response to DAN damage. U2OS cells transfected with Flag-GNL3 WT or the SIM mutant were treated with control or 20 μM ETO for 2 hours and assayed by PLA using anti-Flag and anti-BLM antibodies. Shown are representative confocal images at low magnification. **(c)** BLM SIM is critical for its interaction with GNL3 in response to DNA damage. U2OS cells transfected with Flag-BLM WT or the SIMdm mutant were treated with control or 20 μM ETO for 2 hours and assayed by PLA using anti-Flag and anti-GNL3 antibodies. Shown are representative confocal images at low magnification. **(d)** U2OS cells transfected with control, Flag-BLM or Flag-BLM$^{2KR}$ mutant were treated with control or 20 μM ETO for 2 hours and assayed by PLA using anti-Flag and anti-GNL3 antibodies. Shown are representative confocal images at low magnification. **(e)** BLM SUMOylation is critical for its interaction with GNL3 in response to DNA damage. U2OS cells transfected with GFP, GFP-BLM or GFP-BLM$^{2KR}$ mutant were treated with control or 20 μM ETO for 2 hours and assayed by PLA using anti-GFP and anti-GNL3 antibodies.

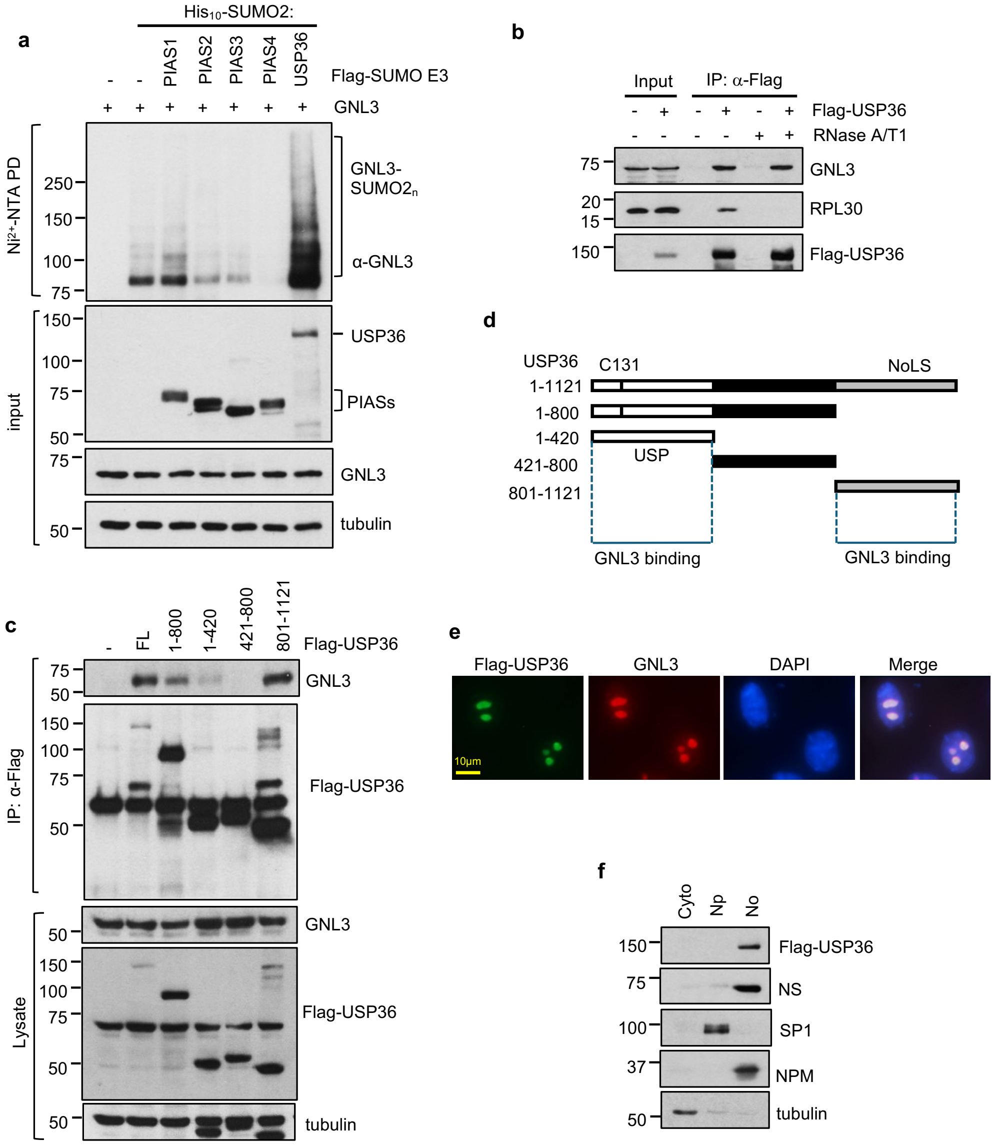


**Figure S6. USP36 is a GNL3 SUMO ligase. (a)** USP36 promotes GNL3 SUMOylation in cells. U2OS-His$_{10}$-SUMO2 cells transfected with indicated plasmids were assayed by $Ni^{2+}$-NTA PD under denaturing conditions followed by IB. The SUMO2 modified GNL3 is indicated. The protein expression is shown at the bottom panels. **(b)** USP36 interacts with GNL3 independently of RNA. 293 cells transfected with Flag-USP36 were assayed by co-IP using anti-Flag antibody in the absence or presence of of 100 µg/mL RNase A and 100U/ml RNase T1 followed by IB. **(c)(d)** Mapping the GNL3 binding to USP36. H1299 cells transfected with Flag-USP36 or the indicated its deletion mutants were assayed by Co-IP using anti-Flag antibody followed by IB (c). Diagram of USP36 deletion mutants is shown in (d). **(e)**. USP36 colocalizes with GNL3 in the nucleolus. U2OS cells transfected with Flag-USP36 were subjected to IF staining with anti-Flag and anti-GNL3 antibodies. **(f)** HeLa cells stably expressing Flag-USP36 were fractionated into cytoplasmic (Cyto), nucleoplasmic (Np) and nucleolar fractions (No) followed by IB. Tubulin, SP1 and nucleophosmin (NPM) were used as cytoplasmic, nucleoplasmic, and nucleolar markers, respectively.

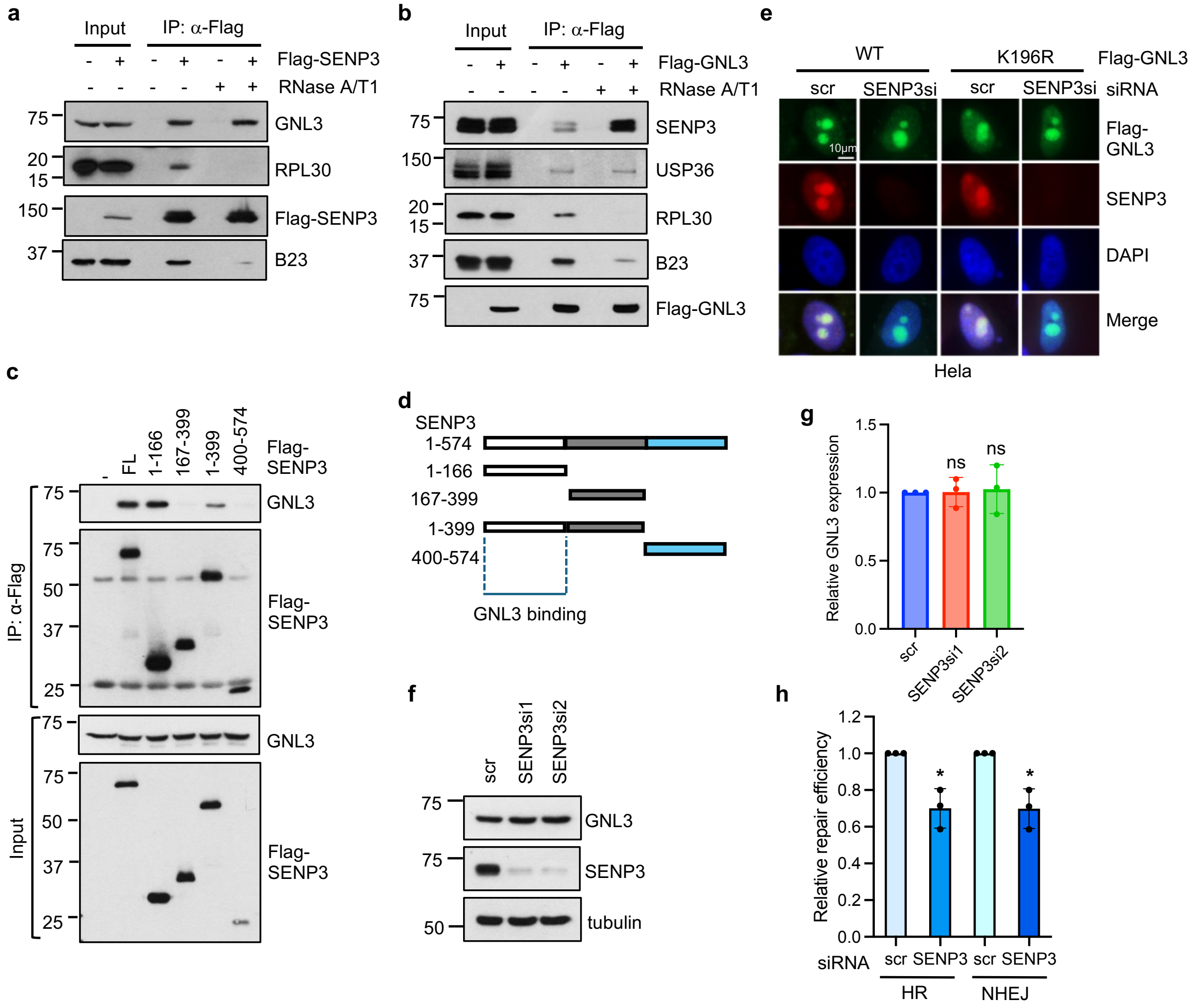


**Figure S7. SENP3 deSUMOylates GNL3. (a)(b)** SENP3 interacts with GNL3 independently of RNA. 293 cells transfected with Flag-SENP3(a) or Flag-GNL3 (b) were assayed by co-IP using anti-Flag antibody in the absence or presence of of 100 μg/mL RNase A and 100U/ml RNase T1 followed by IB. **(c)(d)** The N-terminal SENP3 binds to GNL3. H1299 cells transfected with Flag-SENP3 and its deletion mutants were assayed by co-IP using anti-Flag antibody followed by IB (d). Diagram of SENP3 deletion mutants is shown in (d). (**e**) SENP3 knockdown does not affect GNL3 nucleolar localization. U2OS cells were transfected with Flag-GNL3 (WT or K196R) together with scr or SENP3 siRNA were assayed by IF staining with anti-SENP3 and anti-Flag antibodies. **(f)(g)**. Knockdown of SENP3 does not significantly affect the levels of GNL3. U2OS cells transfected with scr or SENP3 siRNAs were assayed by IB. Shown are the representative IB blots (f) and quantification from three independent experiments (g). (**h**) Knockdown of SENP3 reduces the efficiency of both HR and NHEJ repair. U2OS cells transfected with scr or SENP3 siRNA together with HR or NHEJ reporter plasmids were assayed by flow cytometry for the percentage of GFP-positive cells. Shown are the relative reduction of repair efficiency in SENP3 siRNA transfected cells compared to scr control (*P<0.05).

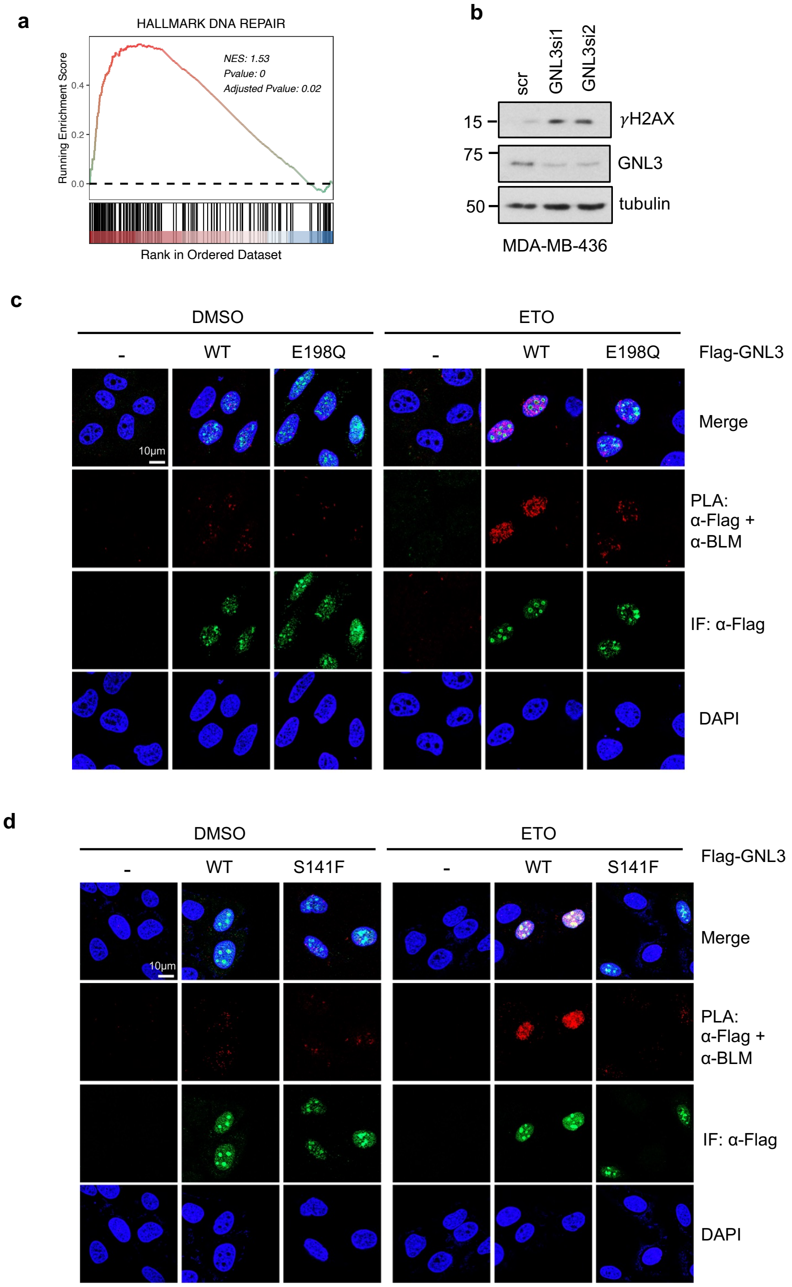


**Figure 8. GNL3 is implicated in breast cancer. (a)** GNL3 expression is associated with DNA damage hallmark in TNBC. Shown is the GSEA score. **(b)** Knockdown of GNL3 induces the levels of γH2AX in MDA-MB-436 cells. **(c)** The GNL3 E198Q mutant showed attenuated interaction with BLM in response to DNA damage. U2OS cells transfected with Flag-GNL3 WT or the E198Q mutant were treated with control or 20 μM ETO for 2 hours and assayed by PLA using anti-Flag and anti-BLM antibodies. Shown are representative confocal images at low magnification. **(d)** The GNL3 S141F mutant showed attenuated interaction with BLM in response to DNA damage. U2OS cells transfected with Flag-GNL3 WT or the S141F mutant were treated with control or 20 μM ETO for 2 hours and assayed by PLA using anti-Flag and anti-BLM antibodies. Shown are representative confocal images at low magnification.

**Supplementary Table 1**

| Plasmid Name | Source | Identifier |
|---|---|---|
| pcDNA3-2Flag-GNL3 | this study | N/A |
| pcDNA3-2Flag-GNL3_K196R | this study | N/A |
| pcDNA3-2Flag-GNL3_K275R | this study | N/A |
| pcDNA3-2Flag-GNL3_G1dm | Dai et al [1] | N/A |
| pcDNA4-TO-2Flag-GNL3 | this study | N/A |
| pcDNA4-TO-2Flag-GNL3/si-1Res | this study | N/A |
| pcDNA4-TO-2Flag-GNL3/K196R/si-1Res | this study | N/A |
| pcDNA4-TO-GNL3/si-1Res | this study | N/A |
| pcDNA4-TO-GNL3_K196R/si-1Res | this study | N/A |
| pcDNA4-TO-2Flag-GNL3_siR_E198Q | this study | N/A |
| pcDNA4-TO-2Flag-GNL3_siR_SIMm | this study | N/A |
| pcDNA4-TO-2Flag-GNL3_siR_S141A | this study | N/A |
| pcDNA4-TO-2Flag-GNL3_siR_S141F | this study | N/A |
| plpcx-2Flag-GNL3 siR | this study | N/A |
| plpcx-2Flag-GNL3 K196R siR | this study | N/A |
| pcDNA3-V5-GNL3 | this study | N/A |
| EGFP-C1-BLM | Addgene | Cat No: 80070 |
| EGFP-C1-BLM DM | Addgene | Cat No: 80071 |
| pCDNA3-2Flag-BLM/1-447 | this study | N/A |
| pCDNA3-2Flag-BLM/1-637 | this study | N/A |
| pcDNA3-2Flag-BLM (FL) | this study | N/A |
| pcDNA3-2Flag-BLM_K317A;K331A | this study | N/A |
| pcDNA3-2Flag-BLM_SIM_dm | this study | N/A |
| pcDNA3-2Flag-Ubc9 | this study | N/A |
| pcDNA3-2Flag-Ubc9 DN | this study | N/A |
| pcDNA3-2Flag-USP36 | Sun et al [2] | N/A |
| pcDNA3-2Flag-USP36/C131A | Sun et al[2] | N/A |
| pcDNA3-2Flag-USP36 1-800 | Sun et al[2] | N/A |
| pcDNA3-2Flag-USP36/1-421 | Sun et al[2] | N/A |
| pcDNA3-2Flag-USP36/801-1121 | Sun et al[2] | N/A |
| pcDNA3-2Flag-USP36/421-800 | Sun et al[2] | N/A |
| pcDNA3-V5-USP36 | Sun et al[2] | N/A |
| pcDNA3-His-SUMO1 | Chen et al[3] | N/A |
| pcDNA3-His-SUMO2 | Sun et al[2] | N/A |
| pcDNA3-His10-SUMO2 | this study | N/A |
| pcDNA3-V5-SUMO2G | this study | N/A |
| pcDNA3-His-Ub | Dai et al[4] | N/A |

| | | |
|---|---|---|
| pcDNA3-2Flag-SENP3 | this study | N/A |
| pcDNA3-2Flag-SENP3/C532S | this study | N/A |
| pcDNA3-2Flag-SENP3/1-166 | this study | N/A |
| pcDNA3-2Flag-SENP3/166-399 | this study | N/A |
| pcDNA3-2Flag-SENP3/1-399 | this study | N/A |
| pcDNA3-2Flag-SENP3/400-574 | this study | N/A |
| pDRGFP | Addgene | Cat No: 26457 |
| pimEJ5GFP | Addgene | Cat No: 44026 |
| pCherry | Dr. Zhenkun Lou[5] | N/A |
| pCBAScel | Addgene | Cat No: 26477 |

**Supplementary Table 2**

| Antibodies | Source | Identifier |
|---|---|---|
| Anti-Flag (IF 1:1000; PLA 1:1000; IB 1:2000 | Sigma | F3165 |
| Anti-V5 (IB 1:3000) | Life technologies | R960-25 |
| Anti-tubulin (IB 1:10000) | Proteintech | 66240-1-Ig |
| Anti-γH2AX (IF 1:500; PLA 1:500; IB 1:4000) | Millipore | 05-636-25 |
| Anti-GNL3 (IF 1:200; PLA 1:200) | Santa Cruz | sc-166460 |
| Anti-GNL3 (IF 1:400; PLA 1:400; IB 1:1000) | Proteintech | 15060-AP |
| Anti-GNL3 (IB 1:2000) | Dai et al | N/A |
| Anti-RAD51 (IF 1:1000) | Abcam | ab133534 |
| Anti-RAD51 (IB 1:2000) | Proteintech | No1496-AP |
| Anti-GAPDH (IB 1:10000) | Proteintech | 60004-1-1g |
| Anti-Nop58 (IF 1:400) | Bethyl | A302-719A |
| Anti-SENP3 (IF 1:400; IB 1:1000) | Cell signaling technology | 5591S |
| Anti-BRCA1 (IF 1:300) | Santa Cruz | sc5654 |
| Anti-BARD1 (IF 1:200) | Bethyl | A300-263A |
| Anti-53BP1 (IF 1:200) | Millipore | 630916 |
| Anti-MDC1 (IF 1:500) | Bethyl | A300-053A |
| Anti-BLM (IF 1:50; IB 1:500) | Santa Cruz | sc-365753 |
| Anti-BLM (PLA 1:200; IB 1:1000) | Invitrogen | PA5-77880 |
| Anti-RPA2 phospho S33 (IF 1:1000; IB 1:5000) | Abcam | ab21187 |
| anti-DNA2 (IB 1:5000) | Proteintech | 18727-1-AP |
| Anti-H2B (IB 1:5000) | Millipore | 07-371 |
| Anti-RPL5 (IB 1:2000) | Dai et al | N/A |
| Anti-RPL30 (IB 1:500) | Santa Cruz | sc-98106 |
| Anti-B23 (IB 1:5000) | Zymed | 32-5200 |
| Anti-SP1 (IB 1:2000) | Millipore | 07-645 |
| Anti-BrdU (IF 1:100; PLA 1:100) | Roche | 11170376001 |
| Anti-SUMO2/3 (IB 1:4000) | Dr. Yoshiaki Azuma | N/A |
| Anti-SUMO2/3 (IB 1:2000) | abcam | ab3742 |
| Anti-USP36 (IB 1:1000) | Dr. Masayuki Komada | N/A |
| Anti-USP36 (IB 1:1000) | Proteintech | 14783-1-AP |
| Alexa Fluor 488 goat anti-mouse IgG (IF 1:400) | Invitrogen | A11001 |
| Alexa Fluor 555 goat anti-mouse IgG (IF 1:400) | Invitrogen | A21422 |
| Alexa Fluor 488 goat anti-rabbit IgG (IF 1:400) | Invitrogen | O6381 |
| Alexa Fluor 555 goat anti-rabbit IgG (IF 1:400) | Invitrogen | A21428 |

### References


1 Dai, M. S., Sun, X. X. & Lu, H. Aberrant expression of nucleostemin activates p53 and induces cell cycle arrest via inhibition of MDM2. *Mol Cell Biol* **28**, 4365-4376 (2008).

2 Sun, X. X. *et al.* The nucleolar ubiquitin-specific protease USP36 deubiquitinates and stabilizes c-Myc. *Proc Natl Acad Sci U S A* **112**, 3734-3739 (2015).

3 Chen, Y. *et al.* The ubiquitin-specific protease USP36 SUMOylates EXOSC10 and promotes the nucleolar RNA exosome function in rRNA processing. *Nucleic Acids Res* (2023).

4 Dai, M. S. *et al.* Ribosomal protein L23 activates p53 by inhibiting MDM2 function in response to ribosomal perturbation but not to translation inhibition. *Mol Cell Biol* **24**, 7654-7668 (2004).

5 Zhu, Q. *et al.* RNF19A-mediated ubiquitination of BARD1 prevents BRCA1/BARD1-dependent homologous recombination. *Nat Commun* **12**, 6653 (2021).
